# Meiotic recombination rate in *Drosophila melanogaster* is robust to hypoxic conditions

**DOI:** 10.64898/2026.11.26.768981

**Authors:** Natalia Rivera-Rincón, Spencer A. Koury, Melika Ghasemi Shiran, Laurie S. Stevison

## Abstract

Meiotic recombination is often described as a plastic trait that responds to environmental stress, but the evidence across stressors is mixed and the role of oxygen availability in particular remains poorly resolved. We tested whether chronic, mild hypoxia alters the rate of meiotic recombination on the X chromosome of *Drosophila melanogaster*. F_1_ females were reared from oviposition through eclosion under one of three conditions: an outside-chamber normoxic control 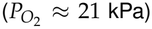, an in-chamber normoxic control 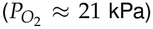, and a hypoxic treatment 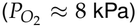. To separate genuine meiotic effects from the confounding influence of differential survival, we paired a six-point multiply-marked test cross with marker-free crosses on identical outcrossed backgrounds, directly measured F_1_ fecundity and F_2_ egg-to-adult viability, and fit the resulting phenotypic class counts to an extended *Cx(Co)*^*m*^ model that accounts for asymmetric meiosis and treatment, marker, and sex-specific mortality. Ovary RNA-seq from unmated F_1_ females showed broad transcriptional changes in the Chamber 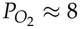 kPa treatment relative to the outside-chamber control, part of which was shared with the in-chamber 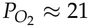 kPa treatment. We saw changes consistent with reduced oxygen, which included up-regulation of stress and DNA damage response genes and down-regulation of ribosome and oxidative phosphorylation pathways. The HIF pathway genes *tango* and *fatiga* were modestly up-regulated relative to the outside-chamber control, but the glycolysis pathway as a whole, which is up-regulated under acute hypoxia, was not. Reduced 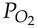 also lowered F_1_ fecundity relative to the in-chamber 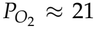 kPa treatment. Despite these physiological and transcriptional signatures of stress, we detected no effect of oxygen treatment on total X chromosome genetic length or on the strength of crossover interference, and no consistent effect on any of the five intervals examined. Recombination on the X chromosome of outbred *D. melanogaster* therefore appears robust to mild chronic environmental hypoxia, even when the same exposure is sufficient to alter ovarian gene expression and depress reproductive output. We suggest that hypoxia-induced plasticity in tracheal growth may buffer ovarian tissue from cellular oxygen limitation, and that the framework used here, combining marker-free viability controls with a process based meiotic model, offers a cleaner way to ask whether other stressors or genotypes produce real recombination plasticity rather than mortality artifacts.

## Introduction

Meiotic recombination is the fundamental property of sexually reproducing organisms where homologous chromosomes exchange genetic material, thereby generating novel allele combinations (reviewed in Stapley et al. 2017). By reshuffling genetic variation each generation, recombination plays a central role in shaping evolutionary trajectories, facilitating adaptation to changing environments, purging of deleterious mutations, and contributing to long-term population persistence (Felsenstein 1974; reviewed in Otto and Lenormand 2002). Recombination rates also exhibit phenotypic plasticity, with early work in *Drosophila melanogaster* demonstrating that temperature treatment can perturb meiotic processes (Plough 1917, 1921; Graubard 1934). Subsequent studies expanded these observations to acute thermal stress (Jackson et al. 2015), nutrient limitations (Neel 1941; Novak et al. 2026) and mating conditions (Zhong and Priest 2010), leading some investigators to propose that recombination rate plasticity is a general feature of organismal stress response to environmental perturbation (Parsons 1988; Stevison *et al*. 2017; Rybnikov *et al*. 2023). However, experimental evidence across environmental stressors remains mixed. For example, no significant changes in mean recombination rates were detected following dietary enrichment with sugar or yeast in *D. melanogaster* (Mostoufi and Singh 2022), or exposure to the dietary flavonoid quercetin (Schramm *et al*. 1998). Similarly, studies examining oxidative stress in honey bees (Langberg *et al*. 2018), and infection in house mice (Dumont *et al*. 2015), found no consistent overall increase in recombination. In contrast, desiccation stress has been associated with genotype dependent changes in recombination rates and crossover interference in *D. melanogaster* (Aggarwal *et al*. 2019). Together, these findings suggest that recombination plasticity is not a universal response to environmental stress, but may instead depend on the nature of the stressor, genetic background, and genomic region. Further-more, recombination rate plasticity under ecologically relevant environmental challenges, such as oxygen availability, remains poorly characterized.

Oxygen availability shapes insect physiology, development, and fitness (Harrison et al. 2018). Harrison *et al*. 2018 define functional hypoxia as “the conditions under which an individual has a demand for *O*_2_ that exceeds its capacity for supply despite compensations”. Here we distinguish environmental hypoxia, which is a reduction in ambient 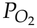, from functional hypoxia in this sense, which also depends on whether physiological compensation matches oxygen supply to demand. Exposure to reduced oxygen levels (such as those found in decaying organic substrates, dense larval aggregations, and microhabitats with limited gas exchange) has profound effects on *D. melanogaster* metabolism (Zhou et al. 2008; Van Voorhies 2009), tracheal re-modeling (Jarecki *et al*. 1999; Lavista-Llanos *et al*. 2002; Centanin et al. 2005), developmental timing (Peck and Maddrell 2005), lifespan (Polan et al. 2020), and reproductive performance (Rivera-Rincón et al. 2026a). The extensive transcriptional and metabolic reprogramming is mediated by an evolutionarily conserved Hypoxia-Inducible Factor (HIF-1*α*) signaling pathway (Bruick and McKnight 2001; Lavista-Llanos et al. 2002; Gorr *et al*. 2006; Romero et al. 2007; Azad et al. 2009, 2012). Under normoxic conditions proline residues of cytosolic HIF-1*α* are hydroxylated by oxygen-dependent Prolyl Hydroxylases (PHD) targeting HIF-1*α* for degradation by proteasomes via binding with the tumor suppressor von Hippel Landau E3 ubiquitin ligase (VHL); however, insufficient levels of oxygen lead to accumulation of HIF-1*α* which then heterodimerizes with oxygen-insensitive HIF-1*β* to act as a transcription factor (reviewed in Semenza 2011). Once the HIF-1*α*/*β* dimer enters the nucleus, it binds to Hypoxia Response Elements (HREs) throughout the genome thereby activating a transcriptional response that includes cellular signaling (Hippo, Notch, JAK-STAT, etc.), stress response (NF*κ*B, mTor, etc.) and causes major shifts in energy metabolism away from oxidative phosphorylation and toward glycolysis (Liu et al. 2006; Zhou et al. 2008; Ding et al. 2022).

Despite strong evolutionary conservation of HIF-1*α* oxygen-sensing pathway, knock-out experiments reveal that the glycolytic response to hypoxia is HIF-independent, at least in *D. melanogaster* third instar larvae (Li et al. 2013; Barretto et al. 2020), and part of the transcriptional response to hypoxia, including metabolic genes, is also HIF-1-independent in *Caenorhabditis elegans* third stage larvae (Shen et al. 2005). Instead, hypoxia transcriptional responses are, at least in part, dependent on Fork-head Box-O signaling (Barretto et al. 2020; Ding et al. 2022) and nuclear Estrogen Related Receptor (Tennessen et al. 2011; Li *et al*. 2013) which are known regulators of metabolism in fly development. Developmental stages differ markedly in their hypoxic exposure and anoxia tolerance (Callier et al. 2015), and population level metabolic responses to hypoxia are further modulated by group interactions and sex (Burggren et al. 2017). Low oxygen concentrations reduce adult and cell size while increasing developmental time (Peck and Maddrell 2005), and early life hypoxia alters adult physiology, reducing lifespan and starvation resistance (Polan et al. 2020). Interestingly, the HIF-1*α* signaling pathway also controls a developmental plastic response in open circulatory system of *D. melanogaster* where tissue specific increased tracheal growth occurs to compensate for low partial pressures of *O*_2_ in a process analogous to angiogenesis in hypoxic tumor tissues of animals with closed circulatory systems (Jarecki et al. 1999; Lavista-Llanos et al. 2002; Centanin et al. 2005). Genomic analysis of experimentally evolved populations identi-fied tracheal growth as the target of hypoxia-based selection in *D. melanogaster* as well as other genetic signals shared with high altitude human populations, underscoring both divergent and convergent nature of adaptation to hypoxic conditions (Jha *et al*. 2015).

Although the physiological and developmental consequences of hypoxia have been extensively studied in *D. melanogaster*, its effects on meiotic recombination remain comparatively poorly understood. Previous studies suggest that oxygen availability can influence recombination, although the magnitude and direction of these responses may depend on genetic background, genomic region, and experimental conditions. Multigenerational selection under chronic hypoxia has been associated with altered recombination rates (Aggarwal *et al*. 2015), whereas more recent single-generation experiments have found no consistent overall effect of hypoxia on X chromosome genetic length, despite evidence of genotype and interval-specific responses and inter-actions with additional environmental stressors (Rivera-Rincón *et al*. 2026b). Together, these findings suggest that hypoxia does not induce a uniform recombination response and highlight the need for experimental approaches that distinguish changes in meiotic recombination from indirect effects of environmental stress on reproductive output and offspring survival. As we noted before, oxygen availability is a ubiquitous and ecologically relevant variable that has not been investigated for its effects on recombination rate plasticity.

Classically, single generation studies, like the one mentioned above, require multiple generation test crosses to study recombination rate plasticity. A major limitation of traditional inference of recombination events is based on scoring F_2_ genotypes as opposed to directly observing F_1_ meiocytes leading to a substantial information loss and statistical confounding due to experimental mortality (Koury *et al*. 2026). Specifically, because only one of four chromatids per bivalent segregates to the functional egg pole in female meiosis and up to 44% of F_2_ individuals do not survive to adulthood to be scored, even under benign crossing conditions *D. melanogaster* recombination studies are typically based on information from only 10-20% of chromatids that are involved in the meiotic events of interest (Koury *et al*. 2026). The former source of loss (asymmetric meiosis) introduces large uncertainty to estimates, while the latter source (experimental mortality) causes both uncertainty and bias in the observable dataset. Furthermore, if experimental variation in recombination rates is induced by strong environmental or genetic perturbation, additional biases due to chromosome segregation defects and differential survival may act to confound inferences about F_1_ meiosis from comparing observations on F_2_ offspring (e.g. Singh *et al*. 2015; Hatkevich *et al*. 2017). This is particularly important since several studies suggest that variation in recombination rate, including plastic changes, are adaptive (Samuk *et al*. 2020; Rybnikov *et al*. 2023; Ritz *et al*. 2017), but without direct con-current measures of experimental mortality to rule out biased differences, these changes cannot assessed under this proposed framework.

There are three approaches to dealing with these difficulties in recombination test crosses. First, experimental methods such as half-tetrad analysis with compound chromosomes (Beadle and Emerson 1935; Chovnick *et al*. 1970), full-factorial marker arrays (Hayman and Parsons 1962; Koury 2023), and rigorous control of genetic backgrounds (Zwick *et al*. 1999b; Koury 2023) can minimize biases and information loss. Second, direct quantification of viability can define the limits, or possibly exclude,confounding effects of experimental mortality (e.g. Sturtevant and Beadle 1936; Redfield 1957; Koury et al. 2026). Finally, probabilistic modeling of the underlying meiotic processes (with explicit treatment of differential survival) allows sound statistical inference from the biased F_2_ data to the F_1_ meiotic events of interest (e.g. Zwick et al. 1999a; Koury et al. 2026). Here, we incorporate all three approaches to test for hypoxia-induced recombination rate plasticity in *D. melanogaster*.

We employ an experimental and statistical analysis capable of separating the effects of reduced oxygen availability from multiple confounding sources of variation. First, we conducted the recombination testcrosses in controlled, outcrossed genetic backgrounds and incorporated normoxic 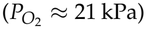 control treatments both inside and outside of airtight environmental chambers to isolate the biological effects of oxygen limitation from chamber related artifacts. Second, because hypoxia imposes stress on female reproductive physiology (Rivera-Rincón *et al*. 2026a) and visible mutant markers cause experiment mortality (Koury et al. 2026), we conducted marker-free control crosses on identical genetic backgrounds to directly quantify F_1_ female fe-cundity and F_2_ egg-to-adult viability without the confounding ef-fect of markers. Third, we analyzed the recombination testcross F_2_ phenotypic class counts with an extended *Cx(Co)*^*m*^ counting model that explicitly accounts for the data-generating process including random, marker-induced, and treatment-induced sources of experimental mortality. This integrative approach allows us to test whether the low oxygen treatment 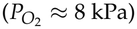 is associated with changes in meiotic recombination while limiting confounding by selective mortality and marker-associated viability differences.

## Materials and methods

### Experimental design

Experimental analysis of *D. melanogaster* X chromosome recombination was conducted in a modified testcross design recently reported in Koury *et al*. (2026). This modified design estimates absolute egg-to-adult viability for a six-point testcross covering spanning 98% of the X chromosome as well as the control viability of a marker-free X chromosome on identical outcrossed genetic backgrounds (Figure 1a, Figure S1a). The seven visible markers of the multiply-marked X chromosome were *yellow, y*^*1*^, *scute, sc*^*1*^, *crossveinless, cv*^*1*^, *vermilion, v*^*1*^, *forked, f*^*1*^, *carnation, car*^*1*^, and *Dp(1;1)sc*^*V1*^, *yellow, y*^*+*^ (Figure 1b) and the four wildtype autosomal backgrounds were Oregon-R (BDSC 25211), Canton-S (BDSC 64349), *w*^*1118*^ (BDSC 6326), and *Drosophila melanogaster* Genetic Reference Panel (DGRP) line 399 (BDSC 25192) (Figure S1b). Please see Koury *et al*. (2026) for full description of stock construction and crossing schemes. All crosses were conducted in polystyrene vials with standard cornmeal *Drosophila* media containing 0.01 v/v McCormick blue food coloring with 50 *µ*L of live yeast paste (0.2 g/mL) and 0.02 v/v blue food coloring (Schneider et al. 2025). Irrespective of treatment, all vials and environmental chambers were housed in a Percival incubator at 21°, 101-2 kPa, 70% relative humidity, and 12:12 hour light-dark cycle.

**Figure 1.**
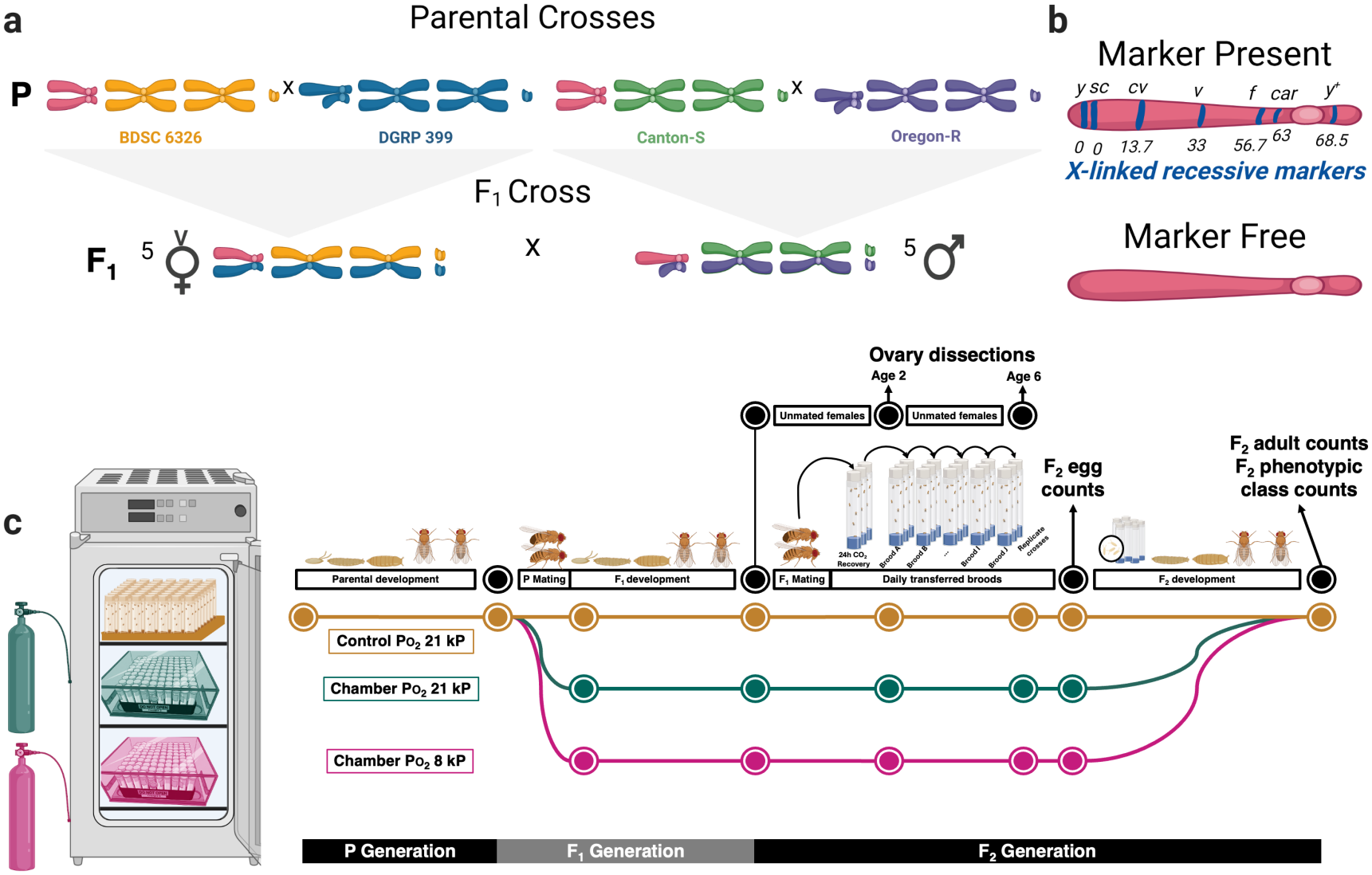
Experimental Design Workflow. **(a)** Here, we used an outcrossing design where two parental crosses were performed to generate heterozygous F_1_ female and male progeny. Chromosomes are depicted schematically to illustrate genotypic composition. First, BDSC 6326 (yellow) was crossed to DGRP 399 (blue). Then, Canton-S (green) was crossed to Oregon-R (purple). The resulting F_1_ progeny of each parental cross were subsequently crossed to each other to generate F_2_ progeny. **(b)** Indicated in pink, each background genome has either an X-chromosome which is marked with X-linked recessive visible markers to count F_2_ phenotypic classes for recombination, or a marker free X-chromosome. **(c)** Two Biospherix chambers were used to control oxygen concentration experimentally, with an outside of chamber control at atmospheric O_2_ concentrations 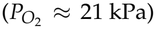. Pre-mixed airgas tanks of either 21% O_2_ (teal) or 8% O_2_ (pink) were used to establish within chamber normoxic conditions 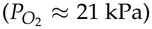 and hypoxic conditions 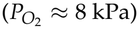. Parental crosses were each placed into one of the three treatment groups with the F_1_ generation developing completely within each treatment condition. All sets of F_1_ crosses underwent 24-hour tap transfers on Drosophila media with blue food dye to measure both F_2_ egg counts and F_2_ adult counts in order to compute egg-to-adult viability measures with and without markers. Additionally, unmated F_1_ females were dissected and their ovaries were used for RNA sequencing. BioRender.org was used to build parts of this figure. License link to be included upon acceptance.

#### Treatment conditions

F_1_ experimental females were generated by parental (P) cross of *DGRP 399 y*^*1*^; *DGRP 399*; *DGRP 399* females to male *y*^*1*^ *sc*^*1*^ *cv*^*1*^ *v*^*1*^ *f*^*1*^ *car*^*1*^ *Dp(1;1)sc*^*V1*^ *y*^*+*^; *BDSC 6326*; *BDSC 6326* (see Figure S1 for crossing scheme). The P crosses were placed under their randomly assigned experimental treatments immediately following mating and were continually maintained under these conditions, thus the F_1_ experimental females experienced the treatment conditions from oviposition through eclosion (Figure 1c). Normoxic (“Chamber 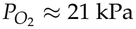”) and hypoxic (“Chamber 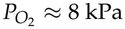”) treatments were established in custom environmental chambers (BioSpherix) flushed with certified gas mixtures (21% O_2_ and 8% O_2_ by volume, respectively, provided by AirGas). Flushing occurred every 12 hours or every time the chambers were opened, to manipulate the oxygen availability following the procedure of Rivera-Rincón *et al*. (2026a). To account for potential chamber-related artifacts, an external normoxic control (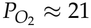 kPa in Auburn, AL referred to as “Control 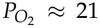 kPa”) was conducted outside of the environmental chambers but still in the same Percival incubator (Figure 1c).

#### Animal husbandry

Following the methods of Koury *et al*. (2026), DGRP 399/BDSC 6326 heterozygous F_1_ experimental females (either with or without markers) were collected over a 72-hour period. After aging for an additional 72-hour period, these females were mated to similarly aged Canton-S/Oregon-R heterozygous males (either with or without markers from a P cross also conducted inside of chambers, Figure 1b) under light CO_2_ anesthesia. Twelve replicate crosses for each treatment were performed at a density of 5 female and 5 males per vial (Figure 1c). Following a 24-hour CO_2_ recovery period, the mated flies were tap transferred onto fresh food every 24 hours for 7 days to generate 7 broods. In total this generated 504 vials of F_2_ eggs (2 marker states*×* 3 treatments *×* 12 replicates *×* 7 broods). Immediately after tap transferring flies, food surface was imaged for egg counting (Schneider et al. 2025). 72 hours after imaging, a 15 cm^2^ piece of Kimwipe was inserted to food and wetted with 100 *µ*L of 0.01M propionic acid 0.001M phosphoric acid.

### F_1_ Female ovarian RNA sequencing

Twenty-four ovary samples (3 treatments*×* 4 replicates *×* 2 time points) were collected for RNA sequencing, each consisting of ten marker-free virgin F_1_ experimental females (Figure S1a) with four biological replicates per treatment (Control 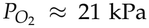, Chamber 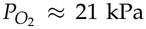, and Chamber 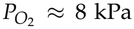) at 2 and 6 days post-eclosion (Figure 1c). All tools and work surfaces were treated with RNase decontamination solution and dissections were performed under RNase-free conditions with flies CO_2_ anesthetized for no longer than ten minutes. Ovaries attached to the oviduct were removed by gently pulling the posterior abdomen while stabilizing the thorax with forceps. Ovary pairs were immediately transferred to pre-chilled 100–200 *µ*L RNAshield (Zymo Research) in 1.5 mL tubes and kept on ice during dissection. Samples were stored at *−*80^*°*^C until RNA extraction.

#### RNA extraction and quantification

RNA was extracted using the Direct-zol RNA Miniprep Plus kit (Zymo Research kit number R2071-A) following the manufacturer’s protocol with minor modifications. RNA concentrations and purity were assessed using a NanoDrop spectrophotometer (Thermo Fisher Scientific), and all samples had absorbance ratios of 260/280 and 260/230 greater than 2. RNA was further purified and concentrated using the RNA Clean & Concentrator-25 kit (Zymo Research kit number R1017) then re-measured using a NanoDrop spec-trophotometer and confirmed with the Qubit RNA BR Assay Kit (Thermo Fisher Scientific). Additional quality checks were done independently by Novogene using Gel Electrophoresis and Agilent 2100 analysis, confirming high quality RNA with all samples exhibiting RNA integrity numbers greater than 9.

#### RNA-seq library preparation and sequencing

RNA library preparation and sequencing were conducted by Novogene. The 24 RNA-seq libraries generated between 12.2 million and 44.1 million raw reads per sample with sequencing quality con-sistently high across all libraries (Q20 values ranging from 98.65% *−*99.28% and Q30 values 95.83% *−*96.84%). Average sequencing error rate was low (approximately 0.01%), GC content was stable, and *≥*97% reads were retained as clean reads. Between 90.81% to 95.44% of clean reads uniquely mapped to the *D. melanogaster* reference genome with pearson correlation demonstrating strong concordance among biological replicates (*R*^2^ *>* 0.94).

#### RNA-seq statistical analysis

Differential expression and enrichment analysis between our comparisons were performed by Novogene. First, the ‘DESeq2’ R package (Love *et al*. 2014) was used to identify differentially expressed genes. The resulting *p*-values were adjusted using the Benjamini and Hochberg’s procedure to control the false discovery rate at 5%. Genes with an adjusted *p*-value < 0.05 were assigned as differentially expressed. Statistical enrichment of differentially expressed genes was then conducted for Gene Ontology (GO) and Kyoto Encyclopedia of Genes and Genomes (KEGG) pathways using the ‘clusterPro-filer’ R package (Yu *et al*. 2012) using genes with an unadjusted *p*-value < 0.05 as input; terms and pathways were considered significantly enriched at a Benjamini–Hochberg adjusted *p* < 0.05. Enrichment results for every pairwise comparison, pooled and for each sampling day, are given in Table S10. To compare the magnitude of pathway-level changes among comparisons, we also summarized DESeq2 log_2_ fold changes (median and interquartile range) across all genes in four KEGG pathways (ribosome, oxidative phosphorylation, glycolysis/gluconeogenesis and the citrate cycle), with pathway membership from the KEGG database (Table S11); no additional tests were performed. All analyses were performed using the reference genome from *D. melanogaster* (BDGP6, Ensembl version GCA_000001215_4). All statistical analyses and visualizations were performed in R version 4.5.3 (R Core Team 2026), with gene-set overlaps visualized using UpSetR (Conway *et al*. 2017) and ‘ggplot2’, and volcano plots and GO/KEGG enrichment plots generated using ‘ggplot2’ (Wickham 2016).

### F_**1**_ Fecundity and F_**2**_ egg-to-adult viability assays

Daily F_1_ female fecundity was measured as the number of eggs per individual per vial from the 24 hour brooding scheme as scored in digital images independently by two research assistants (Schneider et al. 2025; Koury et al. 2026). The degree of undercounting in digital images was assessed on an independent dataset and raw counts were adjusted accordingly (Schneider et al. 2025; Koury et al. 2026) (Figure S2a). Starting 10 days after egg laying and continuing until all viable adults had eclosed, counts of emerging adults were scored for sex (Figure 1c). Egg-to-adult viability of F_2_ individuals was calculated as the number of F_2_ adults eclosing from a vial divided by the mean adjusted F_2_ egg counts.

#### Statistical analysis of fecundity

The identical full factorial experimental design for marker-free and multiply-marked crosses allows F_1_ fecundity data to be analyzed jointly for the effect of visible markers as well as any interaction with treatment, brood, sex, etc. Daily F_1_ individual female fecundity (*F*_*ijkl*_) was modeled as:

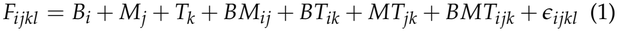

where *B*_*i*_ represents the *i*^*th*^ brood coded as a categorical variable, *M*_*j*_ specifies marker presence or absence in the cross, *T*_*k*_ denotes the treatment, and *ϵ*_*ijkl*_ is the residual error term. This model was analyzed as a 7 *×*2*×* 3 full factorial ANOVA with type III sums of squares in R, using package ‘emmeans’ for post hoc analyses (Lenth and Piaskowski 2025).

#### Statistical analysis of viability

Angular transformation of F_2_ egg-to-adult viability (*V*_*ijklmn*_, the proportion of eggs surviving to adulthood) was analyzed in relation to the same explanatory variables while including the per vial egg density *D*_*i*_ as a covariate (as recommended in Sokal and Rohlf 1995, chapter 13). An additional effect of F_2_ sex as also included (making a 2 *×* 7 *×* 2 *×* 3 design with sex *S*_*j*_, brood *B*_*k*_, marker presence *M*_*l*_, and treatment *T*_*m*_ as fixed effects, categorical variables analyzed with a type III sums of squares). Because embryos cannot be phenotypically sexed, the egg-to-adult viability analysis requires assuming a 1:1 primary sex ratio of all eggs in digital counts. The equal slopes assumption of ANCOVA was violated due to a Density-by-Marker interaction (Figure S2b), so the 2 *×* 7 *×* 2*×* 3 full-factorial design was analyzed as regression with continuous Density *D*_*i*_ and Density-by-Marker *DM*_*il*_ covariates. Thus the ordinary least squares regression model analyzed for F_2_ egg-to-adult viability (*V*_*ijklmn*_) was:

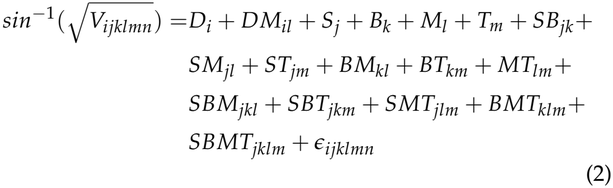

where the residual errors (*ϵ*_*ijklmn*_) are normally distributed with a mean of zero on the transformed scale. Statistical analysis using R packages ‘car’ and ‘emmeans’ were performed on the arcsine transformation of the proportion of eggs surviving to adulthood (*V*_*ijklmn*_), effect estimates and confidence intervals were back-transformed for graphing (Fox and Weisberg 2019; Lenth and Piaskowski 2025).

### Genetic length and crossover interference estimation

Observer effects were not investigated for either fecundity or viability analyses. However, prior to fitting the extended *Cx(Co)*^*m*^ counting model to the multi-locus dataset, differences in single-locus marker recovery rates were calculated from sex-specific F_2_ adult phenotypic classes emerging from multiply-marked crosses for each of the three recombination data collectors. The goodness-of-fit for these recovery rates was tested against the single-locus mendelian expectations assuming all experimental mortality was random with respect to markers and sex (Table S1). Unsurprisingly, this *χ*^2^ test was rejected for all loci and all observers due to mutant viability effects, but the additive property of *χ*^2^ statistics also allows a two degree of freedom *χ*^2^ test of heterogeneity among observers (Table S2). This *χ*^2^ heterogeneity test was also rejected for all loci (Table S2). However, the degree of difference between observers never exceeded 2% deviation in single marker recovery rate, indicating that while there is increased uncertainty because of a statistically significant observer effect, the bias introduced by this effect is relatively small.

#### Fitting the data-generating process

Counts of sex-specific adult phenotypic class for all 19,476 F_2_ adult flies emerging from multiply-marked crosses were recorded, excepting 100 unidentifiable individuals. Additionally, male phenotypic class counts were imputed in a single vial (B15) using the sex-specific, treatment-specific, brood-specific weighted averages scaled to the total egg count of that vial. Finally, five vials with fewer than 10 total individuals eclosing (including two vials with zero eggs) were excluded from analysis. The F_2_ phenotypic class count data from each of the remaining 247 multiply-marked cross vials were fit to an extended *Cx(Co)*^*m*^ counting model of crossover patterning recently described in Koury *et al*. (2026). Briefly, this model assumes that: 1) programmed double strand breaks occur as a homogeneous Poisson point process along the genetic map, 2) crossover maturation in break repair is a stationary renewal process, 3) chromosome segregation is random one-half thinning of this process, 4) fertilization by X-bearing versus Y-bearing sperm is mendelian, and 5) egg-to-adult survival is binomially distributed with a sex-specific, treatment-specific rate parameter determined by the complement of F_2_ marker alleles. Maximum likelihood estimates of model parameters (density of double strand breaks, genetic lengths, and strength of crossover interference) were obtained by downhill simplex method implemented in custom R scripts using the package ‘dfoptim’ to constrain viability estimates to between 0 and 1 (Varadhan et al. 2023) (see Data Availability).

#### Statistical analysis of recombination and crossover interference

After *Cx(Co)*^*m*^ model fitting, the estimated genetic length of the X chromosome from each vial was analyzed as a full-factorial ANOVA (7×3×3 corresponding to seven broods *B*_*i*_, three data collector *O*_*j*_, and three treatments *T*_*k*_ all coded as fixed effects, categorical variables, with type III sums of squares) in the linear model:

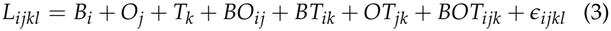

where *ϵ*_*ijkl*_ is the residual error term. Analysis of linear model and pairwise post-hoc tests were performed using R package ‘emmeans’ (Lenth and Piaskowski 2025). Second order meiotic phenotypes (e.g. crossover interference) show high variability when fit to small sample size (i.e. vials as the experimental unit); therefore, strength of crossover interference under each experimental treatment is reported from pooled counts in Table 1, while per-vial estimates of crossover interference and of the five interval lengths were analyzed with the same linear model as total genetic length.

**Table 1.** Summary table for total F_1_ fecundity, mean F_2_ egg-to-adult viability, estimated whole X chromosome genetic lengths, and strength of crossover interference under all experimental designs and experimental treatments.

|  | Marker-free cross |  | Multiply-marked cross |  |  |  |
|---|---|---|---|---|---|---|
|  | F<sub>1</sub> Fecundity | F<sub>2</sub> Survival | F<sub>1</sub> Fecundity | F<sub>2</sub> Survival | X Chr. Length<sup>*a*</sup> | Interference<sup>*b*</sup> |
| Control *P*<sub>*O*<sub>2</sub></sub> ≈ 21 kPa | 13109 | 0.90 | 8642 | 0.66 | 78.5 | 2 |
| Chamber *P*<sub>*O*<sub>2</sub></sub> ≈ 21 kPa | 18307 | 0.84 | 11400 | 0.67 | 80.9 | 1 |
| Chamber *P*<sub>*O*<sub>2</sub></sub> ≈ 8 kPa | 16288 | 0.88 | 9707 | 0.63 | 78.0 | 2 |

<sup>*a*</sup> genetic lengths in centiMorgans estimated from full datasets fit to the extended *Cx(Co)*<sup>*m*</sup> counting model
<sup>*b*</sup> crossover interference parameter *m* estimated from full datasets fit to the extended *Cx(Co)*<sup>*m*</sup> counting model

## Results

Summing across all treatments and both cross designs, 360 F_1_ experimental females produced 77,453 eggs with 60,859 F_2_ adults scored (Table 1). Additionally, 240 marker-free F_1_ females were dissected for ovary gene expression and 19,476 F_2_ adults from multiply-marked crosses were scored for X chromosome recombination.

### Ovary transcriptional responses to hypoxic conditions

Comparison of gene expression in marker-free F_1_ female ovaries pooled across sampling time points revealed 1,079 differentially expressed genes (DEGs) in at least one of the three pairwise treatment comparisons (Table S8). Of these, nearly all (95%; N=1022/1079) were between the Control 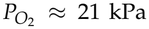 and Chamber 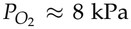 treatment (Figure 2a; Figure 2c; Table S8). In enrichment analyses (see Methods), Chamber 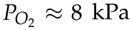 ovaries showed, relative to the Control, down-regulation of the ribosome and oxidative phosphorylation KEGG path-ways (Figure 2b; Table S10) and of ribosomal, mitochondrial and oxidoreductase-activity GO terms (Figure S5), alongside up-regulation of GO terms for cellular response to stress, cellular response to DNA damage stimulus, DNA repair, and several metabolic processes (Figure S4). KEGG pathways for homologous recombination, mismatch repair and DNA replication were also up-regulated (Figure 2b). At the gene level, two of the four core HIF pathway genes (Lavista-Llanos et al. 2002; Centanin et al. 2005; Romero et al. 2007), *tango* and *fatiga*, had modestly higher expression (log_2_ fold change 0.16 and 0.13), whereas *sima* and *Vhl* did not differ (Figure S6; Table S9). Neither *tango* nor *fatiga* differed significantly between the two chamber treatments, although *fatiga* was higher in Chamber 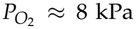 than in Chamber 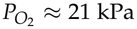 by a similar amount (log2 fold change 0.13), and did not differ between Chamber 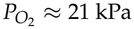 and the Control (Table S9). *Succinate dehydrogenase subunit D (complex II)* had lower expression, and the glycolytic enzyme *pyruvate kinase* had higher expression on Day 6 (Table S9).

**Figure 2.**
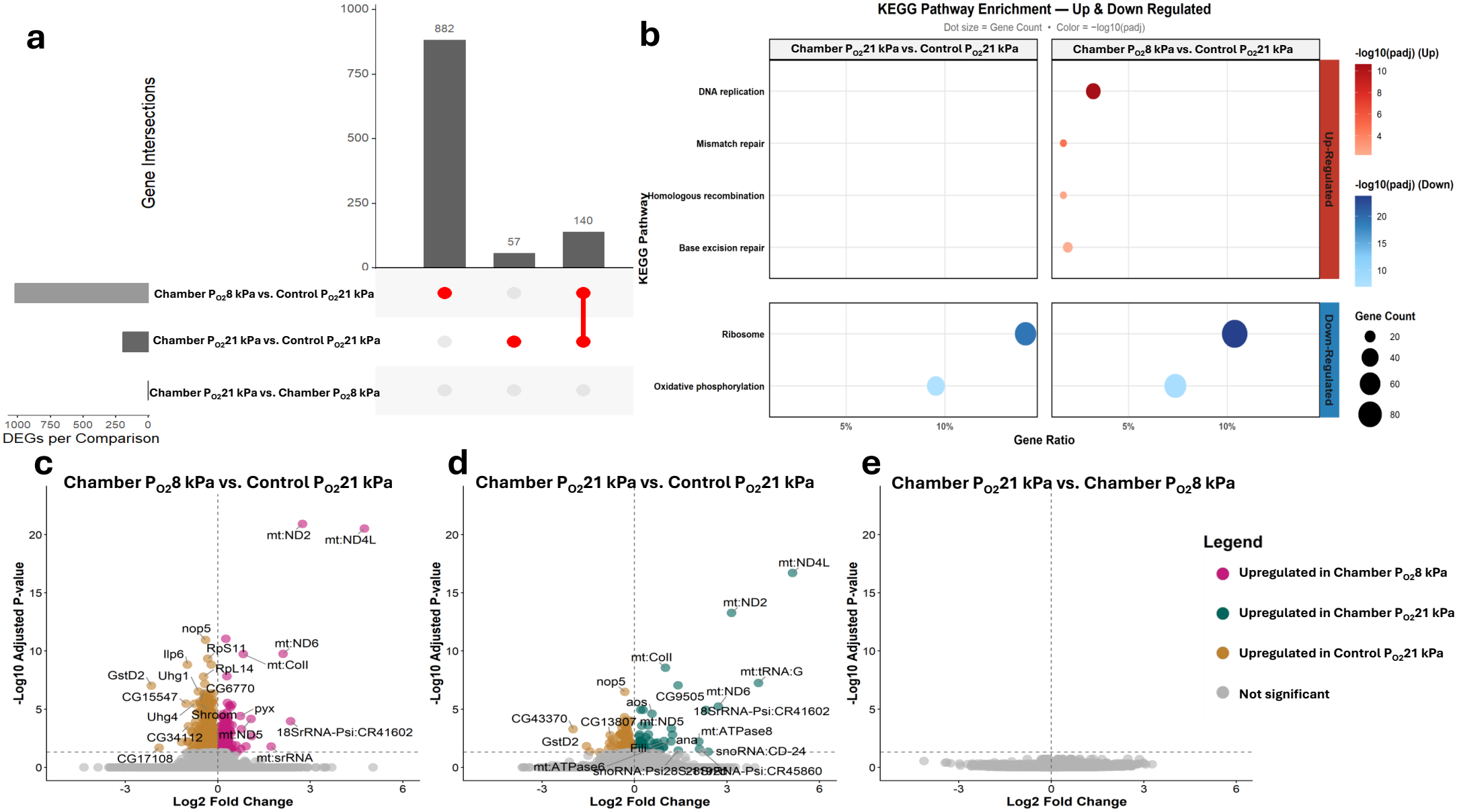
**(a)** UpSet plot showing intersections of significantly differentially expressed genes (DEGs) across three pairwise comparisons: chamber 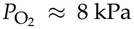 vs. outside-chamber control 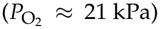, chamber 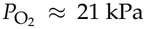 vs. outside-chamber control, and chamber 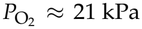 vs. chamber 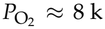Pa. Connected dots indicate DEGs shared between comparisons, while single dots represent DEGs unique to one comparison. Vertical bars show intersection sizes, and horizontal bars show the total number of DEGs per comparison. No significant DEGs were detected between the two chamber treatments when sampling time points were analyzed together. **(b)** KEGG pathway enrichment analysis of upregulated and downregulated genes in each chamber treatment 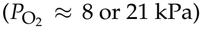 relative to the outside-chamber control 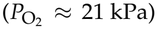. Bubble size represents gene count, and the horizontal axis indicates gene ratio. Red and blue bubbles represent pathways enriched among upregulated and downregulated genes, respectively; color intensity indicates *−* log_10_ of the adjusted *p*-value. **(c–e)** Volcano plots showing differential gene expression for **(c)** chamber 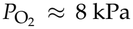 vs. outside-chamber control, **(d)** chamber 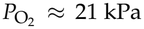 vs. outside-chamber control, and **(e)** chamber 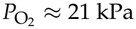 vs. chamber 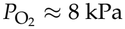. Pink, teal, and gold points indicate genes with significantly higher expression in the chamber 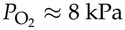 treatment, chamber 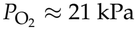 treatment, and outside-chamber control, respectively; gray points indicate genes that did not meet the significance criteria. The vertical axis shows *−*log_10_ of the adjusted *p*-value. Dashed lines indicate significance thresholds, and selected genes are labeled.

Because the chambers differ from the outside Control in more than 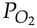(e.g., accumulation of CO_2_ and nitrogenous waste, humidity), we refer to differences between the Chamber 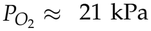 treatment and the Control 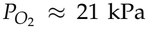 as chamber effects. These accounted for 18% of DEGs (N=197/1,079; Figure 2d), including 57 not shared with any other comparison (Figure 2a). On Day 2, 27% of DEGs (N=466/1,735) were attributable to chamber effects, compared with 14% (N=170/1,222) on Day 6 (Figure S3b, Figure S3e; Table S8). Chamber effects included down-regulation of the ribosome and oxidative phosphorylation pathways also seen in the Chamber 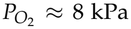 comparison (Figure 2b; Figure S5; Table S10). However, these pathways were reduced more in Chamber 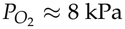 than in Chamber 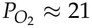 kPa relative to the Control (median log_2_ fold change across pathway genes *−*0.19 vs. *−* 0.11 for the ribosome and *−*0.15 vs. *−*0.05 for oxidative phosphorylation; Table S11).

Pooled across time points, no genes were differentially expressed between the two chamber treatments (Figure 2a). This null result masks a strong effect of time. On Day 2, 82% of DEGs (N=1,426/1,735) distinguished the two chambers, compared with only 5.5% (N=67/1,222) on Day 6 (Figure S3c, Figure S3f; Table S8). Most of these Day 2 differences were unique to the comparison between the two chambers (N=1,081), whereas on Day 6 most DEGs were unique to the Chamber 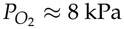 versus Control comparison (N=988; Figure S3g). Conversely, the Chamber 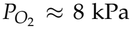 versus Control comparison accounted for 15% of DEGs on Day 2 (N=254/1,735) but 97% on Day 6 (N=1,188/1,222) (Figure S3a, d; Table S8). On Day 2, relative to the Chamber 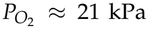, Chamber 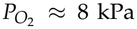 ovaries showed down-regulation of KEGG pathways for carbon metabolism, the citrate (TCA) cycle, glycolysis/gluconeogenesis, amino acid metabolism and the lysosome, together with GO terms for oxidoreductase activity and carbohydrate metabolism (Table S10). Genes in these pathways were modestly lower in the Chamber 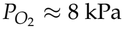 and modestly higher in the Chamber 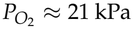 than in the Control, so both treatments contributed to the contrast (Table S11). Conversely, pathways for DNA replication, ATP-dependent chromatin remodeling, and mismatch, nucleotide excision, base excision and homologous recombination repair were up-regulated, along with the GO terms cellular response to stress, cellular response to DNA damage stimulus and DNA repair, paralleling the pooled Chamber 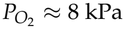 versus Control comparison (Table S10). On Day 6, although only 67 genes differed significantly between the two chambers, the ribosome and oxidative phosphorylation pathways were down-regulated in Chamber 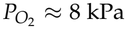 relative to Chamber 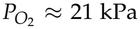 (Table S10). Both pathways were reduced more than twice as much in Chamber 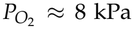 as in Chamber 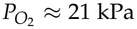 relative to the Control (median log_2_ fold change *−*0.28 vs. *−*0.13 for the ribosome and *−*0.15 vs. *−*0.04 for oxida-tive phosphorylation; Table S11). As a group, glycolysis genes were not up-regulated in any comparison (Table S11), although a few individual genes were, including pyruvate dehydrogenase kinase (*Pdk*), which was higher in Chamber 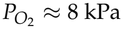 than in Chamber 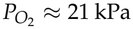 on Day 2 (log_2_ fold change 0.19, adjusted *P* = 0.024).

### Decreased 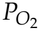 reduces F_1_ fecundity but not F_2_ viability

The statistically significant effect of experimental treatment on daily individual F_1_ female fecundity (*F*_2,457_ = 34.7, *p* = 9.5 *×*10^*−*15^, *η*^2^ = 0.086) was a product of all three treatments dif-fering from each other (Figure 3a, Table 2, Table S3). This main effect can be partitioned into an *increase* of 12 eggs per day per individual due to the chamber effect (*t*_s_(457) = *−*7.98, *p* < 0.001) and a *decrease* of 5 eggs per day per individual due to lowering 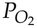 to 8 kPa inside chambers (*t*_s_(457) = *−*3.10, *p* = 0.025). The multiply-marked F_1_ genotype had strongly reduced fecundity relative to the marker-free genotype (*F*_1,457_ = 238, *p* = 1.6 *×*10^*−*43^, *η*^2^ = 0.295), but nonetheless produced a similar pattern of response to experimental treatments (albeit with weaker statistical support, Figure 3a, Table 2, Table S4).

**Figure 3.**
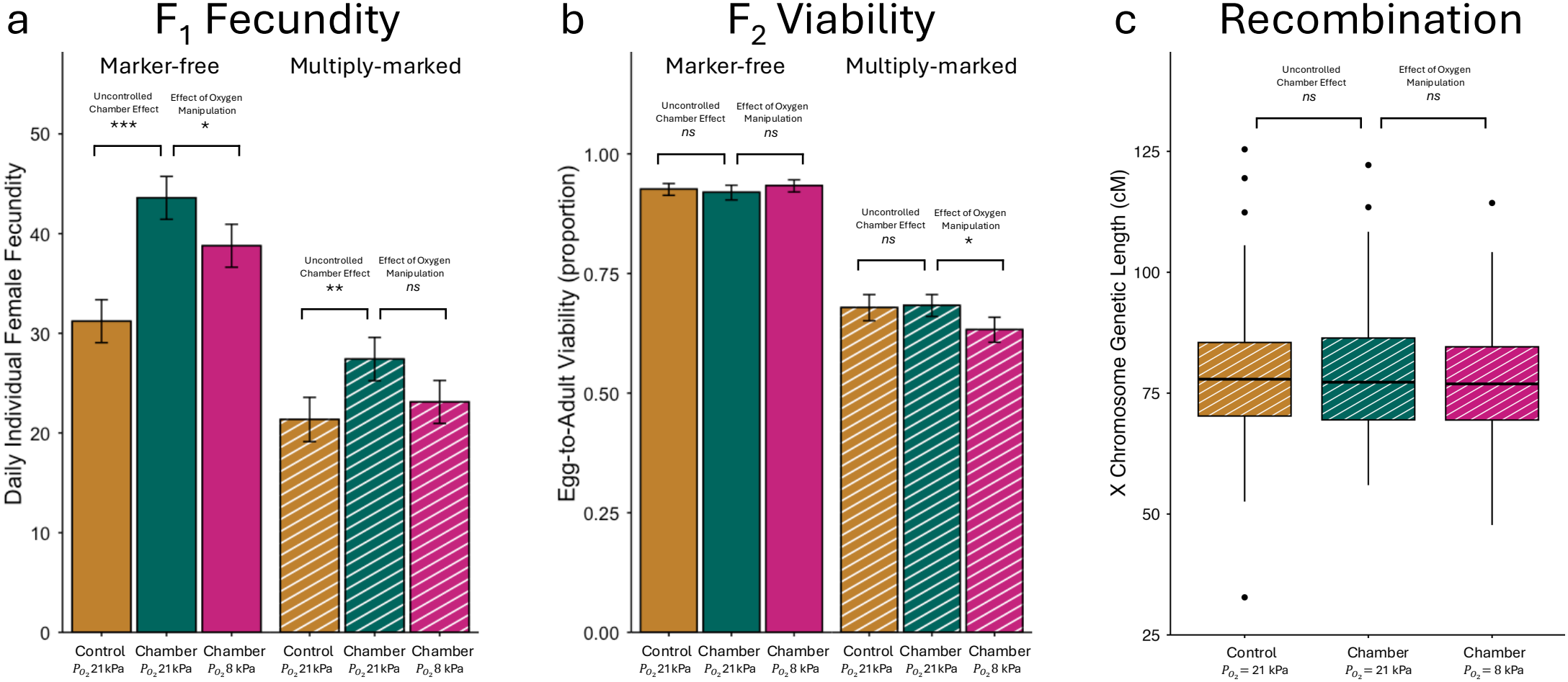
**(a)** Estimated marginal means and 95% confidence intervals for per day per individual F_1_ female fecundity. Marker-free crosses (solid bars) and multiply-marked crosses (white striped bars) depicted for outside the chamber controls at 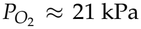 (gold), inside chamber 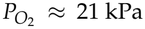 (teal), and inside chamber 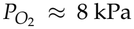 (pink) treatments. **(b)** Estimated marginal means and 95% confidence intervals for F_2_ egg-to-adult viability using the same color-coding scheme (statistical significance assessed on arcsine transformed data but effect estimates and 95% confidence intervals back-transformed for ease of interpretation). **(c)** Boxplots of estimated total X chromosome genetic length in centiMorgans. Each data point is the estimated length from a single vial’s observations fitted to the extended *Cx(Co)*^*m*^ counting model. Statistical significance in all plots based on post-hoc pairwise t-tests of treatment levels (ns: p>0.05, *: 0.05>p>0.01, **: 0.01>p>0.001, ***: 0.001>p after adjusting with Tukey’s method).

**Table 2.** Fecundity: ANOVA table for the per day per individual F_1_ female fecundity under all cross designs and experimental treatments. Statistically significant model effects are presented in bold. Type III sums of squares; the total is the corrected total.

| Source of Variation | *df* | *SS* | *MS* | *F* | *p*-value |
|---|---|---|---|---|---|
| Brood (B) | 6 | 606.5 | 101.1 | 1.00 | 0.42 |
| **Markers (M)** | **1** | **24048** | **24048** | **238** | **1.6×10<sup>-43</sup>** |
| **Treatment (T)** | **2** | **7008** | **3504** | **34.7** | **9.5×10<sup>-15</sup>** |
| B*M Interaction | 6 | 1015 | 169.1 | 1.67 | 0.13 |
| B*T Interaction | 12 | 1416 | 118.0 | 1.17 | 0.30 |
| **M\*T Interaction** | **2** | **1013** | **506.7** | **5.01** | **7.0×10<sup>-3</sup>** |
| B*M*T Interaction | 12 | 486.8 | 40.57 | 0.40 | 0.96 |
| Error | 457 | 46180 | 101.05 |  |  |
| Total | 498 | 81471 |  |  |  |

There was a strong linear dependence of F_2_ egg-to-adult viability on the per vial egg density in marker-free crosses (Figure S2b), but no effect of experimental treatment after accounting for the difference in egg-density (*F*_2,912_ = 1.092, *p* = 0.34) (Figure 3b, Table 3, Table S5). The presence of visible markers decreases survival by 30%, a statistically significant large effect size (*F*_1,912_ = 285, *p* = 7.6*×* 10^*−*56^, *η*^2^ = 0.221) that also over-whelmed any signal of covariation with per vial egg density (Table 3, Figure S2b; Table S6). Post-hoc comparisons reveal that the 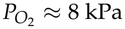 chamber treatment caused greater experimental mortality and was responsible for the small, but significant, Marker-by-Treatment interaction in Table 3 (*F*_2,912_ = 6.107, *p* = 2.3 *×*10^*−*3^, *η*^2^ = 0.0095). Similarly to the fecundity analysis, all statistically significant higher order interactions have small effect sizes (each explaining less than 5% of variation) and are always artifacts of differences involving marker-free versus multiply-marked crosses.

**Table 3.** Viability: Regression analysis for sex-specific, arcsine transformed F_2_ egg-to-adult viability under both cross designs and all experimental treatments with per vial egg density modeled as a covariate. ^1^ Statistically significant model effects are presented in bold.

| Source of Variation | *df* | *SS* | *MS* | *F* | *p*-value |
|---|---|---|---|---|---|
| **Density** | **1** | **0.948** | **0.948** | **39.08** | **6.2×10<sup>-10</sup>** |
| **Density\*Markers** | **1** | **1.050** | **1.050** | **43.27** | **8.0×10<sup>-11</sup>** |
| Sex (S) | 1 | 0.066 | 0.066 | 2.736 | 0.10 |
| Brood (B) | 6 | 0.147 | 0.024 | 1.007 | 0.42 |
| **Markers (M)** | **1** | **6.912** | **6.912** | **285.0** | **7.6×10<sup>-56</sup>** |
| Treatment (T) | 2 | 0.053 | 0.026 | 1.092 | 0.34 |
| S*B Interaction | 6 | 0.104 | 0.017 | 0.711 | 0.64 |
| **S\*M Interaction** | **2** | **1.305** | **1.305** | **53.80** | **4.9×10<sup>-13</sup>** |
| S*T Interaction | 2 | 0.046 | 0.023 | 0.944 | 0.39 |
| **B\*M Interaction** | **6** | **0.403** | **0.067** | **2.768** | **1.1×10<sup>-2</sup>** |
| B*T Interaction | 12 | 0.109 | 0.009 | 0.374 | 0.97 |
| **M\*T Interaction** | **2** | **0.296** | **0.148** | **6.107** | **2.3×10<sup>-3</sup>** |
| **S\*B\*M Interaction** | **6** | **0.371** | **0.062** | **2.550** | **1.8×10<sup>-2</sup>** |
| S*B*T Interaction | 12 | 0.093 | 0.008 | 0.319 | 0.97 |
| S*M*T Interaction | 2 | 0.004 | 0.002 | 0.085 | 0.92 |
| B*M*T Interaction | 12 | 0.296 | 0.025 | 1.017 | 0.43 |
| S*B*M*T Interaction | 12 | 0.260 | 0.022 | 0.891 | 0.56 |
| Error | 912 | 22.12 | 0.024 |  |  |
| Total | 997 | 31.24 |  |  |  |

### Neither chambers effects nor 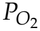 alter recombination

Statistical analysis of the total X chromosome genetic lengths estimated from fitting extended *Cx(Co)*^*m*^ counting model reveals no statistically significant effect of experimental treatment (*F*_2,184_ = 0.88, *p* = 0.42) with 68.3% of total variation in length unexplained. When genetic length of the five intervals are analyzed separately (but not independently because of fitted effect of crossover interference makes interval length estimates inter-dependent), again there was no statistically significant effect of experimental treatments (Table S7c–g). Similarly, there was no statistically significant effect of experimental treatment on crossover interference (*F*_2,184_ = 0.63, *p* = 0.53; Table S7b) with 61.9% of total variation in interference unexplained.

Observer effects were detected for both total genetic length (*F*_2,184_ = 8.23, *p* = 3.8*×* 10^*−*4^; Table 4) and crossover interference (*F*_2,184_ = 24.16, *p* = 4.8*×* 10^*−*10^), with an Observer-by-Treatment interaction for crossover interference (*F*_4,184_ = 4.79, *p* = 1.1*×* 10^*−*3^) but not for total genetic length (*p* = 0.31). Among intervals, Observer effects were strongest in those flank-ing *crossveinless* and *carnation*, consistent with differences in scoring these markers. Pairwise post-hoc comparisons of Treatment effects accounting for observer bias show no statistically significant differences among Treatments (Table S7a,b). Low repeatability of recombination rate estimates (68.3% of variation unexplained in genetic map length and 61.9% unexplained in crossover interference) suggests that while there is some observer bias in data collection, each observer scored similar numbers of vials in every treatment, so this bias does not confound comparisons among treatments.

**Table 4.** Recombination: ANOVA table for X chromosome total genetic length in multiply-marked crosses under all three experimental treatments. Statistically significant model effects presented in bold. Type III sums of squares; the total is the corrected total.

| Source of Variation | *df* | *SS* | *MS* | *F* | *p*-value |
|---|---|---|---|---|---|
| Brood (B) | 6 | 1558 | 260 | 1.53 | 0.17 |
| **Observer (O)** | **2** | **2799** | **1400** | **8.23** | **3.8×10<sup>-4</sup>** |
| Treatment (T) | 2 | 298 | 149 | 0.88 | 0.42 |
| B*O Interaction | 12 | 1811 | 151 | 0.89 | 0.56 |
| B*T Interaction | 12 | 3312 | 276 | 1.62 | 0.088 |
| O*T Interaction | 4 | 821 | 205 | 1.21 | 0.31 |
| **B\*O\*T Interaction** | **24** | **6436** | **268** | **1.58** | **0.050** |
| Error | 184 | 31293 | 170 |  |  |
| Total | 246 | 45298 |  |  |  |

## Discussion

Here, reducing 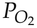 to about 8 kPa altered ovarian gene expression (Figure 2a,c,d; Figure S3) and reduced fecundity (Figure 3a) in outbred, wildtype *D. melanogaster*. The transcriptional signal shows down-regulation of oxidative phosphorylation and ribosome pathways (Figure 2b), which is part of the FOXO-dependent *D. melanogaster* response to hypoxia, which also reduces female fecundity (Ding *et al*. 2022). Similarly, there is up-regulation of cellular stress response, DNA repair, chromatin remodeling pathways, and DNA damage response GO terms (Figure S4) that are also consistent with hypoxia (Mossman *et al*. 2017; Osley et al. 2007). The Chamber 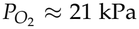 treatment shared part of this signal, including down-regulation of ribo-some and oxidative phosphorylation pathways, but with far fewer DEGs (Figure 2d; Table S11). Because the chambers differ from the Control in more than oxygen, these shared changes cannot be attributed to oxygen alone (see below). Changes spe-cific to oxygen, seen in the direct comparison between chambers, included up-regulation of stress and DNA repair genes on Day 2 and down-regulation of ribosome and oxidative phosphorylation genes on Day 6, but not the glycolytic shift typical of acute hypoxia (Tables S10 and S11). The modest up-regulation of *tango* and *fatiga* was significant only relative to the Control, although *fatiga* was higher in Chamber 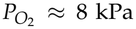 than in Chamber 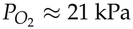 by a similar amount. Despite these physiological and transcriptional effects, we did not see any plastic changes in recombination, suggesting that meiosis in outbred *D. melanogaster* genotypes is robust to mild hypoxic environmental conditions.

This result is consistent with both hypoxia-induced development plasticity of the *D. melanogaster* tracheal system that compensates for reduced O_2_ availability, as well as several recombination rate studies in this genetic model organism that have not found evidence of plasticity across a range of environmental perturbations (Plough 1917; Murray and Lewis 1957; Hinton 1963; Schramm et al. 1998; Jarecki et al. 1999; Lavista-Llanos et al. 2002; Centanin et al. 2005; Mostoufi and Singh 2022). We speculate that the chronic, mild reduction in 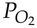 applied throughout larval stages induced developmental plasticity in tracheal system growth, that is similar to angiogenesis in hypoxic tissues of tumors in organisms with closed respiratory systems. Thus while the ambient partial pressure of O_2_ was reduced in the chambers, increased gas exchange through a the more extensive tracheal system may have limited functional hypoxia in ovarian tissue (Harrison *et al*. 2018), consistent with the absence of a canonical cellular hypoxia response and with no observable change in meiotic recombination. This robustness may also reflect the life history of *D. melanogaster*, whose larvae develop in rotting, microbe-rich food where local oxygen levels are low (Callier *et al*. 2015; Lee *et al*. 2019). The species is exceptionally tolerant of acute severe hypoxia, surviving 6 hours at 0.5% O_2_ (Liu *et al*. 2006), and even survives short-term anoxia (Chen *et al*. 2002; Haddad 2006).

Notably, the transcriptional response to Chamber 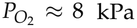 included up-regulation of the homologous recombination pathway (Figure 2b), the repair machinery that also processes meiotic double-strand breaks into crossovers. Differentially expressed genes in this pathway included *Blm, rad50, okra* (Rad54), *mus101, mms4, Pol32* and *RpA-70*, each with 15–25% higher expression than in the Control (Table S9). In *D. melanogaster*, Blm helicase promotes meiotic crossover patterning (Hatkevich *et al*. 2017), yet we detected no change in total genetic length, interval lengths or crossover interference. These expression changes were modest, measured at the mRNA level, and from whole ovaries, in which meiotic cells are a small fraction of the so-matic and germline cells present. Additionally, many of the same genes also function in DNA replication, which was similarly up-regulated. Much like the hypoxia results, this transcriptional response in the recombination machinery did not translate into altered crossover rates or a change in the distribution of

### Direct viability estimates improve recombination plasticity comparisons

In a prior study directly examining the effects of hypoxia on recombination in *D. melanogaster* is the experimental evolution study of Aggarwal *et al*. (2015), which reported altered recom-bination rates after multi-generational selection under chronic low oxygen. However, that design confounds evolutionary responses with within-generation plastic effects, making it difficult to isolate the contribution of hypoxia per se to meiotic recombination. Rivera-Rincón *et al*. (2026a) demonstrated that mild, chronic hypoxia 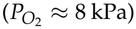 imposes substantial physiological and reproductive stress on *D. melanogaster* females across five genetic backgrounds.

Specifically, hypoxia significantly reduced F_1_ fertility and altered body mass in *D. melanogaster* females, with genotype-dependent effects on thermal tolerance, confirming that the oxygen treatment used here is a genuine stressor at the organismal level. In a companion study to the one cited above, using quadruple X-linked markers, Rivera-Rincón *et al*. (2026b) found that hypoxia alone did not significantly affect total X chromosome genetic map length, with an average decrease of only 0.96 cM. Although the magnitude and direction of the response varied among genotypes, the overall effect was small. In post-hoc comparisons, one strain, DGRP 508, had a significant increase of 6.35 cM in the 3-4 day brood. Additionally, the same strain also increased recombination rate in interval *y-cv* in brood A and another strain DGRP 57 had lower recombination in interval *cv-v*. However, that study was done at a different temperature than was used here as well as different genetic backgrounds. Specifically, they used a backcrossing procedure as opposed to our outcrossing design, only scored male F_2_ adults, and did not quantify experimental mortality. They also compared only within-chamber treatments, thus lacking the external normoxic control included in the present design. Though it is worth noting that they removed adult F_1_ females from the chambers at eclosion, earlier than was done here. Given our transcriptional results show the two chamber treatments being highly differentiated until day 2, this suggests those comparisons did track differences in oxygen exposure. Together, these comparisons suggest that the effects of environmental stress on recombination are context-dependent and that the presence of a substantial physiological response does not necessarily predict a corresponding change in recombination rate.

A critical caveat when comparing these findings with other plasticity studies is that our experimental approach used controlled, outcrossed genetic backgrounds, directly measured egg-to-adult mortality, and modeled the data generating process with the extended *Cx(Co)*^*m*^ counting model (Koury et al. 2026), which explicitly accounts for asymmetric meiosis as well as marker-induced and treatment-induced experimental mortality. This probabilistic framework provides a more robust estimate of genetic map length than the conventional practice of computing recombinant fractions from observed F_2_ phenotypic class frequencies, which is not based on a model of meiosis and can confound meiotic and post-meiotic processes. Interestingly, here we find that marker *cv* accounts for 94% of marker-associated mortality in outcrosses, highlighting the potential of marker-associated viability differences to influence recombination estimates when mortality is not explicitly accounted for. Additionally, by including an outside-of-chamber normoxic control, we were able to distinguish the biological effects of direct oxygen manipulation from uncontrolled chamber-related artifacts. The independent validation through marker-free crosses further strengthens this approach by providing direct measures of F_1_ fecundity and F_2_ egg-to-adult viability under each treatment condition.

### Ovarian gene expression under reduced *P*_*O*2_ partly matches acute responses to hypoxia

Most of what is known about the transcriptional response to hypoxia in *D. melanogaster* comes from short exposures of whole animals: 30 to 120 minutes at 6% O_2_ in adults (Mossman *et al*. 2017), 1 to 6 hours at 0.5% O_2_ in adults (Liu *et al*. 2006), 2.5 hours at 1% O_2_ in adults (Azad *et al*. 2009), or 6 hours at 4% O_2_ in larvae (Li *et al*. 2013). In contrast, our F_1_ females experienced a milder reduction in oxygen 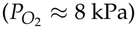 continuously from oviposition through eclosion, and we sampled a single tissue, the ovary, two and six days after eclosion (Figure 1c). Additionally, sensitivity to hypoxia changes across development (Callier *et al*. 2015; Heinrich *et al*. 2011; Stobdan *et al*. 2024), so the transcriptional state we measured likely reflects acclimation to a long-term exposure rather than the initial stress response captured by acute studies.

Several features of our data match the acute literature. *fatiga*, whose *fgaB* isoform is a Sima-dependent hypoxia-inducible gene (Acevedo *et al*. 2010), had higher expression in the Chamber 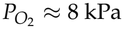 treatment than in the Control (Figure S6). Because gene-level counts pool all isoforms, this modest fold change may underestimate the *fgaB* response. *fatiga* was also higher in Chamber 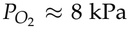 than in Chamber 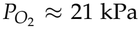 by a similar amount, although not significantly, and did not differ between Chamber 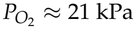 and the Control, so this signal is not explained by chamber effects. Ribosomal protein and oxidative phosphorylation genes were broadly down-regulated, as in the FOXO-dependent response to hypoxia described by Ding *et al*. (2022). *MFS14* and CG31324, both identified as hypoxia-responsive in previous studies (Azad *et al*. 2009; Mossman *et al*. 2017), had higher expression in both chamber treatments than in the Control on Day 6 (Table S9).

However, many genes induced by acute hypoxia had *lower* expression in Chamber 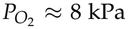 ovaries (Table S9). These included genes regulated by the two main hypoxia-responsive transcription factors. Among HIF targets, *Tribbles*, a direct target of *Sima* (Noguchi *et al*. 2022), was lower than in the Control on Day 6, as was *scylla* (Reiling and Hafen 2004; Liu *et al*. 2006), although *scylla* was higher than in the Chamber 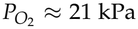 treatment on Day 2. Among FOXO-associated genes, *Thor* (4E-BP), which is up-regulated by acute hypoxia in adults (Liu *et al*. 2006), was lower in every Chamber 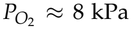 versus Control comparison, and *Hipk*, a FOXO-dependent gene required for hypoxia survival (Ding *et al*. 2022), was also lower. The same pattern held for genes identified in acute screens of adults: CG1600 (Azad *et al*. 2009), *GstD2* (reported as *GstD21* by Liu *et al*. 2006), *astray* (Liu *et al*. 2006), and CG10924 (*Pepck*), which was consistently up-regulated by hypoxia across all female geno-types in Mossman *et al*. (2017). On Day 2, Chamber 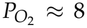 kPa ovaries also had lower expression of *Ldh* (Li *et al*. 2013), *ade5* (Liu *et al*. 2006) and *Mdr49* (Azad *et al*. 2009) than the Chamber 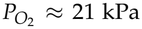 ovaries, along with lower expression of the autophagy gene *Atg8a* and the lysosome pathway (Table S10), even though autophagy is induced by and required for hypoxia tolerance in larvae (Valko *et al*. 2022). Heat shock protein responses were mixed. *Hsp70Aa* and *Hsp70Ab*, which are induced by acute hypoxia (Azad et al. 2009), had higher expression in Chamber 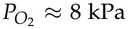 than in Chamber 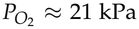 ovaries on Day 2, but *Hsp70Ab* was lower than in the Control on Day 6.

Several explanations, which are not mutually exclusive, could account for these reversals. First, the hypoxia response involves negative feedback: *fatiga*, itself induced by *sima*, hydroxylates *sima* and targets it for degradation (Acevedo et al. 2010), and *scylla* inhibits TOR signaling, of which *thor* is a downstream effector (Reiling and Hafen 2004). Expression of these genes may therefore overshoot and then fall below baseline over a prolonged exposure, depending on where in the feedback cycle the tissue is sampled. Second, the reversals were strongest on Day 6, when the Chamber 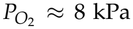 treatment accounted for nearly all DEGs. This timing is consistent with a shift from an initial stress response toward long-term acclimation, including the broad suppression of ribosome biogenesis, oxidative phosphorylation and central carbon metabolism. Third, most reference studies measured whole animals or larvae, whereas the ovary has distinct metabolic and developmental demands. Finally, the transcriptional response to hypoxia in *D. melanogaster* is highly genotype-specific (Mossman et al. 2017), so responses in our single outcrossed genetic background need not match those of the strains used previously.

An alternative interpretation is that Chamber 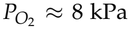 ovaries show a general depression of energy metabolism rather than a cellular response to hypoxia. The response to acute hypoxia in *D. melanogaster* includes up-regulation of glycolysis, which is at least partly HIF-independent (Li et al. 2013). In contrast, glycolysis genes as a group were not up-regulated in any comparison, and on Day 2, genes for glycolysis, the TCA cycle, and amino acid and central carbon metabolism were all lower in Chamber 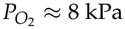 than in Chamber 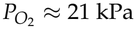 ovaries (Tables S10 and S11). Together with lower ribosome and oxidative phosphorylation expression on Day 6 and reduced fecundity, this pattern is also consistent with metabolic stress, in which reduced energy metabolism lowers reproductive output. The two interpretations are not exclusive, because suppressing metabolic genes is itself a mechanism of hypoxia tolerance in *D. melanogaster* (Zhou et al. 2008). In the terms of Harrison *et al*. (2018), ovaries may have compensated for reduced oxygen supply by lowering demand, so that the treatment imposed environmental hypoxia without sustained functional hypoxia in the tissue. Because we did not measure tissue oxygen or metabolic rate, our data cannot distinguish these possibilities.

Some of these differences also reflect chamber effects rather than oxygen alone. The chambers differed from the outside Control in CO_2_ accumulation, humidity and nitrogenous waste as well as oxygen, and our design cannot distinguish among these factors. Based on published respiration rates, adult flies and the yeast paste in each chamber are expected to consume enough oxygen between flushes to lower 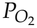, and raise 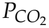, by up to about 0.65 kPa (Supplementary Methods). This is a larger proportional change at 8 kPa than at 21 kPa, so 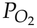 in the low-oxygen chamber likely fell somewhat below 8 kPa between flushes. *GstD2* and CG1600 were lower in both chamber treatments than in the Control on Day 6, *MFS14* was most highly expressed in the Chamber 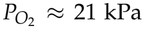 treatment on Day 2, and several metabolic and lysosome genes were higher in the Chamber 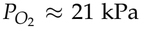 treatment than in the Control on Day 2 (Table S9). The two chamber treatments also converged over time: they differed at 1,426 genes on Day 2 but only 67 on Day 6, while the Chamber 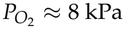 versus Control comparison grew from 254 to 1,188 DEGs (Table S8). This convergence could reflect shared chamber conditions, stage-specific differences in sensitivity to hypoxia (Callier *et al*. 2015; Heinrich *et al*. 2011; Stobdan *et al*. 2024), or both. On Day 2, the lower expression of TCA cycle and glycolytic genes in Chamber 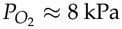 relative to Chamber 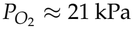 ovaries (Tables S10 and S11) resembles the suppression of these genes by the transcriptional repressor *hairy* in hypoxia-tolerant flies (Zhou *et al*. 2008), although *hairy* itself did not differ. Together, these patterns highlight the value of including multiple experimental controls when interpreting gene expression responses to environmental manipulation.

### Concluding Remarks

Here, we investigated the role of hypoxia to alter meiotic recombination rate. We found that although reduced 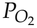 lowered fecundity, as has been shown previously (Rivera-Rincón *et al*. 2026a), meiotic recombination rate was robust to this perturbation. This result is consistent with previous work showing a null result for plasticity in recombination rate. It also suggests that *D. melanogaster* ovaries and subsequent recombination may be particularly tolerant to hypoxia as compared to other perturbations due to their life history. Still, it is worth noting that other recent work has shown temporal changes in recombination rate in other genotypes and specific intervals (Rivera-Rincón *et al*. 2026b). While that study did not use the same process-based method for estimating meiotic recombination rate used here, it is worth noting that genotype-specific responses are common for recombination rate plasticity (Rybnikov *et al*. 2023; Novak *et al*. 2026) as well as for hypoxia (Mossman *et al*. 2017). Having established a methodological framework that explicitly accounts for viability in examining plasticity in recombination, future work can now meaningfully extend this approach to investigate whether the recombination rate robustness observed here holds across other genotypes in the case of hypoxia as well as adapt this methodology to other stressors, such as diet (Novak *et al*. 2026; Mostoufi and Singh 2022). Finally, accounting for differential mortality lends itself well to directly tackling mechanistic explanations for recombination rate plasticity, such as adaptive hypotheses proposed in the literature (Hadany and Beker 2003).

## Data availability

Raw data and code for all analyses have been integrated into a GitHub repository with a static DOI (will request upon acceptance) which can be found at https://github.com/StevisonLab/Hypoxia-Recombination-2026. Also, raw RNA-seq data from ovaries of female *Drosophila melanogaster* exposed to chronic hypoxia have been deposited in the NCBI Gene Expression Omnibus (GEO) under accession number GSE352266.

## Acknowledgments

We would like to thank members of the Stevison Lab for help with the project. Specifically, we thank Cassidy Schneider for help with egg imaging, as well as Connor Haynes and Eva Hammonds for help with egg counts. Claude (Anthropic) and ChatGPT (OpenAI) were used to assist with code generation and debugging for RNA-seq analyses as well as figure preparation. Also Claude (Anthropic) was used during manuscript preparation to audit references against bibliographic databases, to cross-check reported results against outputs, to verify cited claims against source articles, and to assist with drafting and condensing portions of the RNA-seq Results and Discussion.

All analyses were designed and performed by the authors. The authors reviewed and verified all AI-assisted content and take full responsibility for the manuscript.

