## Supplemental Material for "Meiotic recombination rate in *Drosophila melanogaster* is robust to hypoxic conditions"

### Supplementary Methods

#### *Expected change in chamber gas composition*

While our experiment did not directly measure oxygen uptake or oxygen concentration in the chambers, we have roughly calculated the expected $O_2$ in the chambers using published values from the literature. Specifically, each 13.8 L chamber contained 12 vials, each with 10 adults (5 females, 5 males) and 50 µL of 0.2 g/mL baker's yeast. Using a mean adult respiration rate of 5.8 µL $O_2$ mg-1 h-1 and wet masses of 0.69 mg (females) and 0.51 mg (males) (Burggren et al. 2017), adults consume about 10 mL $O_2$ per chamber per day. Using a published respirometry rate for baker's yeast (Simonovik and Gnaiger 2017), the yeast consumes about 170 mL $O_2$ per day. It is worth noting this value comes from measures at 37°C in buffer after rehydration at 30°C, not on fly food at 21°C. With a typical Q10 of about 2, yeast at 21°C would respire roughly 3× less, so the yeast term is probably an overestimate. Together, this is about 180 mL $O_2$ per day, or about 90 mL (0.65% of the chamber volume) in the 12 hours between flushes. This would lower $PO_2$ by about 0.65 kPa before each flush (21 to 20.3 kPa, and 8 to 7.3 kPa) and raise $PCO_2$ by a similar amount. Proportionally, this is a larger change in the low-oxygen chamber (about 8% of available $O_2$) than in the 21 kPa chamber (about 3%). This estimate excludes developing larvae and yeast growth, so it is an approximate, order-of-magnitude estimate.

### Supplementary Tables

**Table S1 Single locus marker recovery rates.** $\chi^2$goodness-of-fit test for the three recombination rate data collectors to mendelian expectations assuming all experimental mortality is random with respect to sex and markers. Loci are a) *scute*, b) *crossveinless*, c) *vermilion*, d) *forked*, e) *carnation*, and f) $Dp(1{:}1)sc^{V1}$, $y^+$.

a

| scute | Observed Count ($O_i$) | | | Expected Count ($E_i$) | | | $((O_i - E_i)^2)/(E_i)$ | | |
|---|---|---|---|---|---|---|---|---|---|
| data collector | MS | NRR | SK | MS | NRR | SK | MS | NRR | SK |
| female_wildtype | 492 | 4309 | 1360 | 569 | 4476 | 1535 | 10 | 6 | 20 |
| male_wildtype | 491 | 3857 | 1281 | 569 | 4476 | 1535 | 11 | 86 | 42 |
| female_mutant | 387 | 2829 | 979 | 569 | 4476 | 1535 | 58 | 606 | 201 |
| male_mutant | 326 | 2317 | 848 | 569 | 4476 | 1535 | 104 | 1041 | 307 |
| Lethal | 876 | 6925 | 2472 | 297 | 2334 | 800 | 1132 | 9030 | 3491 |
| Sum | 2572 | 20237 | 6940 | 2572 | 20237 | 6940 | 1314 | 10769 | 4061 |
| | | | | | | $\chi^2$, df=4, p-value | < 0.001 | < 0.001 | < 0.001 |

b

| Crossveinless | Observed Count ($O_i$) | | | Expected Count ($E_i$) | | | $((O_i - E_i)^2)/(E_i)$ | | |
|---|---|---|---|---|---|---|---|---|---|
| data collector | MS | NRR | SK | MS | NRR | SK | MS | NRR | SK |
| female_wildtype | 511 | 4167 | 1387 | 569 | 4476 | 1535 | 6 | 21 | 14 |
| male_wildtype | 527 | 4038 | 1361 | 569 | 4476 | 1535 | 3 | 43 | 20 |
| female_mutant | 368 | 2971 | 952 | 569 | 4476 | 1535 | 71 | 506 | 221 |

| | | | | | | | | | |
|---|---|---|---|---|---|---|---|---|---|
| male_mutant | 290 | 2136 | 768 | 569 | 4476 | 1535 | 137 | 1223 | 383 |
| Lethal | 876 | 6925 | 2472 | 297 | 2334 | 800 | 1132 | 9030 | 3491 |
| Sum | 2572 | 20237 | 6940 | 2572 | 20237 | 6940 | 1348 | 10823 | 4129 |
| | | | | | | $\chi^2$, df=4, p-value | < 0.001 | < 0.001 | < 0.001 |

c

| Vermilion | Observed Count ($O_i$) | | | Expected Count ($E_i$) | | | $((O_i - E_i)^2)/(E_i)$ | | |
|---|---|---|---|---|---|---|---|---|---|
| data collector | MS | NRR | SK | MS | NRR | SK | MS | NRR | SK |
| female_wildtype | 487 | 4009 | 1303 | 569 | 4476 | 1535 | 12 | 49 | 35 |
| male_wildtype | 505 | 3627 | 1256 | 569 | 4476 | 1535 | 7 | 161 | 51 |
| female_mutant | 392 | 3129 | 1036 | 569 | 4476 | 1535 | 55 | 405 | 162 |
| male_mutant | 312 | 2547 | 873 | 569 | 4476 | 1535 | 116 | 831 | 285 |
| Lethal | 876 | 6925 | 2472 | 297 | 2334 | 800 | 1132 | 9030 | 3491 |
| Sum | 2572 | 20237 | 6940 | 2572 | 20237 | 6940 | 1321 | 10476 | 4024 |
| | | | | | | $\chi^2$, df=4, p-value | < 0.001 | < 0.001 | < 0.001 |

**Table S1 Single locus marker recovery rates (continued).** $\chi^2$ goodness-of-fit test for the three recombination rate data collectors to mendelian expectations assuming all experimental mortality is random with respect to sex and markers. Loci are a) *scute*, b) *crossveinless*, c) *vermilion*, d) *forked*, e) *carnation*, and f) *Dp(1:1)sc$^{V1}$, y$^{+}$*.

d

| Forked | Observed Count ($O_i$) | | | Expected Count ($E_i$) | | | $((O_i - E_i)^2)/(E_i)$ | | |
|---|---|---|---|---|---|---|---|---|---|
| data collector | MS | NRR | SK | MS | NRR | SK | MS | NRR | SK |
| female_wildtype | 488 | 3893 | 1292 | 569 | 4476 | 1535 | 11 | 76 | 38 |
| male_wildtype | 452 | 3430 | 1209 | 569 | 4476 | 1535 | 24 | 244 | 69 |
| female_mutant | 391 | 3245 | 1047 | 569 | 4476 | 1535 | 56 | 338 | 155 |
| male_mutant | 365 | 2744 | 920 | 569 | 4476 | 1535 | 73 | 670 | 246 |
| Lethal | 876 | 6925 | 2472 | 297 | 2334 | 800 | 1132 | 9030 | 3491 |
| Sum | 2572 | 20237 | 6940 | 2572 | 20237 | 6940 | 1296 | 10359 | 4000 |
| | | | | | | $\chi^2$, df=4, p-value | < 0.001 | < 0.001 | < 0.001 |

e

| Carnation | Observed Count ($O_i$) | | | Expected Count ($E_i$) | | | $((O_i - E_i)^2)/(E_i)$ | | |
|---|---|---|---|---|---|---|---|---|---|
| data collector | MS | NRR | SK | MS | NRR | SK | MS | NRR | SK |
| female_wildtype | 464 | 3905 | 1295 | 569 | 4476 | 1535 | 19 | 73 | 37 |
| male_wildtype | 410 | 3335 | 1176 | 569 | 4476 | 1535 | 44 | 291 | 84 |
| female_mutant | 415 | 3233 | 1044 | 569 | 4476 | 1535 | 42 | 345 | 157 |
| male_mutant | 407 | 2839 | 953 | 569 | 4476 | 1535 | 46 | 599 | 221 |
| Lethal | 876 | 6925 | 2472 | 297 | 2334 | 800 | 1132 | 9030 | 3491 |
| Sum | 2572 | 20237 | 6940 | 2572 | 20237 | 6940 | 1283 | 10337 | 3990 |
| | | | | | | $\chi^2$, df=4, p-value | < 0.001 | < 0.001 | < 0.001 |

f

| Dp(1:1)sc$^{V1}$, y$^{+}$ | Observed Count ($O_i$) | | | Expected Count ($E_i$) | | | $((O_i - E_i)^2)/(E_i)$ | | |
|---|---|---|---|---|---|---|---|---|---|
| data collector | MS | NRR | SK | MS | NRR | SK | MS | NRR | SK |
| female_wildtype | 485 | 3909 | 1290 | 569 | 4476 | 1535 | 12 | 72 | 39 |
| male_wildtype | 428 | 3408 | 1184 | 569 | 4476 | 1535 | 35 | 255 | 80 |
| female_mutant | 394 | 3229 | 1049 | 569 | 4476 | 1535 | 54 | 347 | 154 |
| male_mutant | 389 | 2766 | 945 | 569 | 4476 | 1535 | 57 | 653 | 227 |
| Lethal | 876 | 6925 | 2472 | 297 | 2334 | 800 | 1132 | 9030 | 3491 |
| Sum | 2572 | 20237 | 6940 | 2572 | 20237 | 6940 | 1289 | 10357 | 3991 |
| | | | | | | $\chi^2$, df=4, p-value | < 0.001 | < 0.001 | < 0.001 |

**Table S2 Single locus heterogeneity test.** $\chi^2$heterogeneity test for deviation of three data collectors from mendelian expectations. Loci are a) *scute*, b) *crossveinless*, c) *vermilion*, d) *forked*, e) *carnation*, and f) *Dp(1:1)sc$^{V1}$, y$^{+}$*.

a

| Scute | $O_i$ | $E_i$ | $(O_i- Ei)^2/E_i$ | |
|---|---|---|---|---|
| data collector | Pooled | Pooled | Pooled | |
| female_wildtype | 6161 | 6579 | 27 | |
| male_wildtype | 5629 | 6579 | 137 | |
| female_mutant | 4195 | 6579 | 864 | |
| male_mutant | 3491 | 6579 | 1450 | |
| Lethal | 10273 | 3431 | 13643 | |
| Sum | 29749 | 29749 | 16121 | $\chi^2$ goodness-of-fit from pooled counts |
| | | | 16145 | $\Sigma\ \chi^2$ goodness-of-fit from table S1a |
| | | | 24 | $\chi^2$ heterogeneity test, df=2 |
| | | | < 0.001 | p-value heterogeneity test |

b

| crossveinless | $O_i$ | $E_i$ | $(O_i- Ei)^2/E_i$ | |
|---|---|---|---|---|
| data collector | Pooled | Pooled | Pooled | |
| female_wildtype | 6065 | 6579 | 40 | |
| male_wildtype | 5926 | 6579 | 65 | |
| female_mutant | 4291 | 6579 | 796 | |
| male_mutant | 3194 | 6579 | 1742 | |
| lethal | 10273 | 3431 | 13643 | |
| sum | 29749 | 29749 | 16286 | $\chi^2$ goodness-of-fit from pooled counts |
| | | | 16301 | $\Sigma\ \chi^2$ goodness-of-fit from table S1b |
| | | | 14 | $\chi^2$ heterogeneity test, df=2 |
| | | | 0.003 | p-value heterogeneity test |

c

| vermilion | $O_i$ | $E_i$ | $(O_i- Ei)^2/E_i$ | |
|---|---|---|---|---|
| data collector | Pooled | Pooled | Pooled | |
| female_wildtype | 5799 | 6579 | 93 | |
| male_wildtype | 5388 | 6579 | 216 | |
| female_mutant | 4557 | 6579 | 622 | |
| male_mutant | 3732 | 6579 | 1232 | |
| lethal | 10273 | 3431 | 13643 | |
| sum | 29749 | 29749 | 15806 | $\chi^2$ goodness-of-fit from pooled counts |
| | | | 15822 | $\Sigma\ \chi^2$ goodness-of-fit from table S1c |
| | | | 16 | $\chi^2$ heterogeneity test, df=2 |
| | | | < 0.001 | p-value heterogeneity test |

**Table S2 Single locus heterogeneity test (continued).** $\chi^2$heterogeneity test for deviation of three data collectors from mendelian expectations.Loci are a) *scute*, b) *crossveinless*, c) *vermilion*, d) *forked*, e) *carnation*, and f) *Dp(1:1)sc^{V1}, y^+*.

d

| forked | $O_i$ | $E_i$ | $(O_i- Ei)^2/E_i$ | |
|---|---|---|---|---|
| data collector | Pooled | Pooled | Pooled | |
| female_wildtype | 5673 | 6579 | 125 | |
| male_wildtype | 5091 | 6579 | 337 | |
| female_mutant | 4683 | 6579 | 547 | |
| male_mutant | 4029 | 6579 | 989 | |
| lethal | 10273 | 3431 | 13643 | |
| sum | 29749 | 29749 | 15640 | $\chi^2$ goodness-of-fit from pooled counts |
| | | | 15654 | Σ $\chi^2$ goodness-of-fit from table S1d |
| | | | 14 | $\chi^2$ heterogeneity test, df=2 |
| | | | < 0.001 | p-value heterogeneity test |

e

| carnation | $O_i$ | $E_i$ | $(O_i- Ei)^2/E_i$ | |
|---|---|---|---|---|
| data collector | Pooled | Pooled | Pooled | |
| female_wildtype | 5664 | 6579 | 127 | |
| male_wildtype | 4921 | 6579 | 418 | |
| female_mutant | 4692 | 6579 | 541 | |
| male_mutant | 4199 | 6579 | 861 | |
| lethal | 10273 | 3431 | 13643 | |
| sum | 29749 | 29749 | 15591 | $\chi^2$ goodness-of-fit from pooled counts |
| | | | 15610 | Σ $\chi^2$ goodness-of-fit from table S1e |
| | | | 19 | $\chi^2$ heterogeneity test, df=2 |
| | | | < 0.001 | p-value heterogeneity test |

f

| Dp(1:1)$sc^{V1}$, $y^+$ | $O_i$ | $E_i$ | $(O_i- Ei)^2/E_i$ | |
|---|---|---|---|---|
| data collector | Pooled | Pooled | Pooled | |
| female_wildtype | 5684 | 6579 | 122 | |
| male_wildtype | 5020 | 6579 | 370 | |
| female_mutant | 4672 | 6579 | 553 | |
| male_mutant | 4100 | 6579 | 934 | |
| lethal | 10273 | 3431 | 13643 | |
| sum | 29749 | 29749 | 15622 | $\chi^2$ goodness-of-fit from pooled counts |
| | | | 15637 | Σ $\chi^2$ goodness-of-fit from table S1f |
| | | | 15 | $\chi^2$ heterogeneity test, df=2 |
| | | | < 0.001 | p-value heterogeneity test |

#### Table S3 Marker-free $F_1$ fecundity post-hoc analysis of treatment levels

| Experimental design | Treatment contrast | Estimate | SE | df | t.ratio | p.value |
|---|---|---|---|---|---|---|
| Marker-free cross | Control – Chamber 8 kPa | -7.57 | 1.55 | 457 | -4.88 | < 0.001 |
| Marker-free cross | Control – Chamber 21 kPa | -12.38 | 1.55 | 457 | -7.98 | < 0.001 |
| Marker-free cross | Chamber 8 kPa - Chamber 21 kPa | -4.81 | 1.55 | 457 | -3.10 | 0.025 |

#### Table S4 Multiply-marked $F_1$ fecundity post-hoc analysis of treatment levels

| Experimental design | Treatment contrast | Estimate | SE | df | t.ratio | p.value |
|---|---|---|---|---|---|---|
| Multiply-marked cross | Control – Chamber 8 kPa | -1.76 | 1.57 | 457 | -1.12 | 0.875 |
| Multiply-marked cross | Control – Chamber 21 kPa | -6.06 | 1.58 | 457 | -3.84 | 0.002 |
| Multiply-marked cross | Chamber 8 kPa - Chamber 21 kPa | -4.30 | 1.56 | 457 | -2.77 | 0.065 |

#### Table S5 Marker-free $F_2$ egg-to-adult survival post-hoc analysis of treatment levels

| Experimental design | Treatment contrast | Estimate | SE | df | t.ratio | p.value |
|---|---|---|---|---|---|---|
| Marker-free cross | Control – Chamber 8 kPa | -0.015 | 0.018 | 912 | -0.826 | 0.963 |
| Marker-free cross | Control – Chamber 21 kPa | 0.012 | 0.019 | 912 | 0.648 | 0.987 |
| Marker-free cross | Chamber 8 kPa - Chamber 21 kPa | 0.027 | 0.017 | 912 | 1.548 | 0.633 |

#### Table S6 Multiply-marked $F_2$ egg-to-adult survival post-hoc analysis of treatment levels

| Experimental design | Treatment contrast | Estimate | SE | df | t.ratio | p.value |
|---|---|---|---|---|---|---|
| Multiply-marked cross | Control – Chamber 8 kPa | 0.049 | 0.017 | 912 | 2.810 | 0.057 |
| Multiply-marked cross | Control – Chamber 21 kPa | -0.005 | 0.018 | 912 | -0.266 | 1.000 |
| Multiply-marked cross | Chamber 8 kPa - Chamber 21 kPa | -0.053 | 0.017 | 912 | -3.065 | 0.027 |

**Table S7 Recombination rate and interference post-hoc analysis of treatment levels.**

a Total Map Length

| Treatment contrast | Estimate | SE | df | t.ratio | p.value |
|---|---|---|---|---|---|
| Control – Chamber 8 kPa | -3.37 | 2.81 | 184 | -1.20 | 0.46 |
| Control – Chamber 21 kPa | -3.06 | 2.81 | 184 | -1.09 | 0.52 |
| Chamber 8 kPa - Chamber 21 kPa | 0.31 | 2.81 | 184 | 0.11 | 0.99 |

b Interference

| Treatment contrast | Estimate | SE | df | t.ratio | p.value |
|---|---|---|---|---|---|
| Control – Chamber 8 kPa | -0.08 | 0.65 | 184 | -0.12 | 0.99 |
| Control – Chamber 21 kPa | 0.59 | 0.65 | 184 | 0.90 | 0.64 |
| Chamber 8 kPa - Chamber 21 kPa | 0.66 | 0.65 | 184 | 1.03 | 0.56 |

c Interval 1 (*yellow–crossveinless*) [model x5]

| Treatment contrast | Estimate | SE | df | t.ratio | p.value |
|---|---|---|---|---|---|
| Control – Chamber 8 kPa | -1.35 | 1.41 | 184 | -0.96 | 0.61 |
| Control – Chamber 21 kPa | -1.33 | 1.41 | 184 | -0.94 | 0.61 |
| Chamber 8 kPa - Chamber 21 kPa | 0.02 | 1.41 | 184 | 0.01 | 1.00 |

d Interval 2 (*crossveinless–vermilion*) [model x4]

| Treatment contrast | Estimate | SE | df | t.ratio | p.value |
|---|---|---|---|---|---|
| Control – Chamber 8 kPa | -0.84 | 1.41 | 184 | -0.60 | 0.82 |
| Control – Chamber 21 kPa | -0.54 | 1.41 | 184 | -0.38 | 0.92 |
| Chamber 8 kPa - Chamber 21 kPa | 0.30 | 1.40 | 184 | 0.21 | 0.98 |

e Interval 3 (*vermilion–forked*) [model x3]

| Treatment contrast | Estimate | SE | df | t.ratio | p.value |
|---|---|---|---|---|---|
| Control – Chamber 8 kPa | -2.63 | 1.48 | 184 | -1.78 | 0.18 |
| Control – Chamber 21 kPa | -3.39 | 1.48 | 184 | -2.30 | 0.06 |
| Chamber 8 kPa - Chamber 21 kPa | -0.76 | 1.47 | 184 | -0.51 | 0.86 |

f Interval 4 (*forked–carnation*) [model x2]

| Treatment contrast | Estimate | SE | df | t.ratio | p.value |
|---|---|---|---|---|---|
| Control – Chamber 8 kPa | -0.08 | 1.23 | 184 | -0.07 | 1.00 |
| Control – Chamber 21 kPa | 0.96 | 1.23 | 184 | 0.78 | 0.72 |
| Chamber 8 kPa - Chamber 21 kPa | 1.04 | 1.23 | 184 | 0.85 | 0.68 |

g Interval 5 (*carnation–$y^+$ duplication*) [model x1]

| Treatment contrast | Estimate | SE | df | t.ratio | p.value |
|---|---|---|---|---|---|
| Control – Chamber 8 kPa | 1.53 | 0.78 | 184 | 1.98 | 0.12 |
| Control – Chamber 21 kPa | 1.24 | 0.78 | 184 | 1.60 | 0.25 |
| Chamber 8 kPa - Chamber 21 kPa | -0.29 | 0.77 | 184 | -0.38 | 0.92 |

**Table S8 Summary of differentially expressed genes across the experiment.** Number of differentially expressed genes (DEGs) in each pairwise treatment comparison, with sampling days pooled and for each sampling day separately. DEGs were identified with DESeq2 (Benjamini–Hochberg adjusted $P < 0.05$; no fold-change cutoff). Up and Down give the direction of expression in the first-named treatment. "%" is relative to the number of unique DEGs across the three comparisons within that analysis (pooled, 1,079; Day 2, 1,735; Day 6, 1,222). A gene can be differentially expressed in more than one comparison, so percentages within an analysis sum to more than 100. 8 kPa = Chamber $PO_2 \approx 8$ kPa; 21 kPa = Chamber $PO_2 \approx 21$ kPa; Control = outside-chamber $PO_2 \approx 21$ kPa.

| Analysis | Comparison | Up | Down | Total | Union | %-age |
|---|---|---|---|---|---|---|
| Pooled | Chamber 8 kPa vs Control | 501 | 521 | 1022 | 1079 | 94.7 |
| | Chamber 21 kPa vs Control | 73 | 124 | 197 | 1079 | 18.3 |
| | Chamber 8 kPa vs Chamber 21 kPa | 0 | 0 | 0 | 1079 | 0 |
| Day 2 | Chamber 8 kPa vs Control | 99 | 155 | 254 | 1735 | 14.6 |
| | Chamber 21 kPa vs Control | 201 | 265 | 466 | 1735 | 26.9 |
| | Chamber 8 kPa vs Chamber 21 kPa | 662 | 764 | 1426 | 1735 | 82.2 |
| Day 6 | Chamber 8 kPa vs Control | 514 | 674 | 1188 | 1222 | 97.2 |
| | Chamber 21 kPa vs Control | 33 | 137 | 170 | 1222 | 13.9 |
| | Chamber 8 kPa vs Chamber 21 kPa | 7 | 60 | 67 | 1222 | 5.5 |

**Table S9.** Expression of known hypoxia-responsive genes in $F_1$ ovaries. Expected responses are from the cited studies, most of which used acute exposures of whole adults or larvae. The last seven rows are not known hypoxia-responsive genes, but instead are enriched genes from the KEGG homologous recombination pathway (dme03440) mentioned in text. Values are $\log_2$ fold changes (DESeq2) for the first-named treatment, with Benjamini–Hochberg adjusted P; ns = not significant (padj ≥ 0.05). 8 kPa = Chamber PO2 ≈ 8 kPa; 21 kPa = Chamber PO2 ≈ 21 kPa; Control = outside-chamber PO2 ≈ 21 kPa. Mean FPKM is across all 24 samples.

| Category | Gene | FlyBase ID | Expected response to hypoxia | Reference | Mean FPKM | Pooled: 8 kPa vs Control | Pooled: 21 kPa vs Control | Pooled: 8 kPa vs 21 kPa | Day 2: 8 kPa vs Control | Day 2: 21 kPa vs Control | Day 2: 8 kPa vs 21 kPa | Day 6: 8 kPa vs Control | Day 6: 21 kPa vs Control | Day 6: 8 kPa vs 21 kPa |
|---|---|---|---|---|---|---|---|---|---|---|---|---|---|---|
| **HIF pathway** | *sima* | FBgn0266411 | mRNA unchanged (protein stabilized) | Lavista-Llanos et al. 2002 | 29.8 | ns | ns | ns | ns | ns | ns | ns | ns | ns |
| **HIF pathway** | *tgo* | FBgn0264075 | Constitutive | Romero et al. 2007 | 30.2 | +0.16 (padj=0.017) | ns | ns | ns | ns | ns | ns | ns | ns |
| **HIF pathway** | *Vhl* | FBgn0041174 | Regulates Sima stability | Romero et al. 2007 | 5.1 | ns | ns | ns | ns | ns | ns | ns | ns | ns |
| **HIF pathway** | *fatiga (Hph)* | FBgn0264785 | Up (fgaB isoform) | Acevedo et al. 2010 | 40.8 | +0.13 (padj=0.029) | ns | ns | ns | ns | ns | ns | ns | ns |
| **HIF pathway** | *scylla* | FBgn0041094 | Up | Reiling & Hafen 2004; Liu et al. 2006; Li et al. 2013 | 117.3 | ns | ns | ns | ns | ns | +0.26 (padj=0.0057) | -0.17 (padj=0.026) | ns | ns |
| **HIF pathway** | *Tribbles* | FBgn0028978 | Up (Sima target) | Noguchi et al. 2022; Li et al. 2013 | 22.4 | ns | ns | ns | ns | ns | ns | -0.38 (padj=0.0026) | ns | ns |
| **HIF pathway** | *Ldh (ImpL3)* | FBgn0001258 | Up | Li et al. 2013 | 1.7 | ns | ns | ns | ns | ns | -0.87 (padj=0.037) | ns | ns | ns |
| **HIF pathway** | *Pyk* | FBgn0267385 | Glycolytic enzyme; not hypoxia-induced in wild-type larvae | Li et al. 2013 | 284.2 | ns | ns | ns | ns | ns | ns | +0.16 (padj=0.016) | ns | ns |
| **HIF pathway** | *Musashi* | FBgn0011666 | mRNA unchanged (protein down) | Bertolin et al. 2016 | 9.5 | ns | ns | ns | ns | -0.31 (padj= | +0.22 (padj=0.025) | ns | ns | ns |

| | | | | | | | | | | | | | | |
|---|---|---|---|---|---|---|---|---|---|---|---|---|---|---|
| | | | | | | | | | | 0.004) | | | | |
| **FOXO-associated** | *Thor (4E-BP)* | FBgn0261560 | Up | Liu et al. 2006; Li et al. 2013 | 108 | -0.64 (padj=0.00068) | ns | ns | -0.70 (padj=0.02) | -0.68 (padj=0.00068) | ns | -0.57 (padj=0.0013) | ns | ns |
| **FOXO-associated** | *Hipk* | FBgn0035142 | Up | Ding et al. 2022 | 14.7 | -0.16 (padj=0.00048) | ns | ns | ns | ns | ns | -0.24 (padj=1.4e-05) | -0.20 (padj=0.0033) | ns |
| **FOXO-associated** | *Relish* | FBgn0014018 | Up | Barretto et al. 2020 | 31.8 | ns | ns | ns | ns | ns | ns | ns | ns | ns |
| **Metabolism** | *Pepck (CG10924)* | FBgn0034356 | Up | Mossman et al. 2017 | 3 | -1.17 (padj=0.0067) | ns | ns | ns | ns | -0.75 (padj=0.038) | -1.61 (padj=0.0016) | ns | ns |
| **Metabolism** | *hairy* | FBgn0001168 | Up | Mossman et al. 2017; Zhou et al. 2008 | 11.4 | ns | ns | ns | ns | ns | ns | ns | ns | ns |
| **Metabolism** | *SdhC* | FBgn0037873 | Up in one female background | Mossman et al. 2017 | 55.8 | ns | ns | ns | ns | ns | ns | -0.27 (padj=0.037) | ns | ns |
| **Metabolism** | *SdhD* | FBgn0039112 | Up in one female background | Mossman et al. 2017 | 51.9 | -0.23 (padj=0.0097) | ns | ns | ns | ns | ns | ns | ns | ns |
| **Acute screens** | *MFS14* | FBgn0010651 | Up | Azad et al. 2009; Mossman et al. 2017 | 5 | ns | +0.94 (padj=0.02) | ns | ns | +0.74 (padj=0.0017) | -0.51 (padj=0.029) | +1.06 (padj=2.3e-05) | +1.27 (padj=3.2e-06) | ns |
| **Acute screens** | *CG31324* | FBgn0051324 | Up | Mossman et al. 2017 | 2 | ns | ns | ns | ns | ns | ns | +1.08 (padj=0.014) | +1.31 (padj=0.0073) | ns |
| **Acute screens** | *CG1600* | FBgn0033188 | Up | Azad et al. 2009; Liu et al. 2006 | 24.6 | -0.54 (padj=0.0017) | ns | ns | ns | ns | -0.42 (padj=0.047) | -0.81 (padj=6.5e-08) | -0.80 (padj=1.9e-07) | ns |

| | | | | | | | | | | | | | | |
|---|---|---|---|---|---|---|---|---|---|---|---|---|---|---|
| **Acute screens** | *GstD2 (GstD21)* | FBgn0010038 | Up | Liu et al. 2006 | 5.3 | -2.16 (padj=9.6e-08) | -1.56 (padj=0.015) | ns | ns | ns | -0.92 (padj=0.011) | -3.03 (padj=8.8e-18) | -3.23 (padj=6.9e-20) | ns |
| **Acute screens** | *astray* | FBgn0023129 | Up | Liu et al. 2006 | 69.9 | ns | ns | ns | ns | ns | ns | -0.18 (padj=0.041) | ns | ns |
| **Acute screens** | *ade5* | FBgn0020513 | Up | Liu et al. 2006 | 84.8 | ns | ns | ns | ns | ns | -0.54 (padj=0.00086) | ns | ns | ns |
| **Acute screens** | *Mdr49* | FBgn0004512 | Up | Azad et al. 2009 | 2.5 | ns | ns | ns | ns | ns | -0.74 (padj=0.029) | ns | ns | ns |
| **Stress and autophagy** | *Hsp70Aa* | FBgn0013275 | Up | Azad et al. 2009 | 0.5 | ns | ns | ns | ns | ns | +1.32 (padj=0.038) | ns | ns | ns |
| **Stress and autophagy** | *Hsp70Ab* | FBgn0013276 | Up | Azad et al. 2009 | 1.9 | ns | -0.54 (padj=0.043) | ns | ns | ns | +1.18 (padj=0.014) | -0.84 (padj=0.028) | ns | ns |
| **Stress and autophagy** | *Hsp23* | FBgn0001224 | Up | Azad et al. 2009 | 2.5 | ns | ns | ns | ns | ns | ns | ns | ns | ns |
| **Stress and autophagy** | *Atg8a* | FBgn0052672 | Up | Valko et al. 2022 | 166.3 | ns | ns | ns | -0.14 (padj=0.046) | ns | -0.28 (padj=8.2e-05) | ns | ns | ns |
| **Homologous recombination** | *Blm* | FBgn0002906 | Not hypoxia-related (discussed in text) | KEGG dme03440; Hatkevich et al. 2017 | 24.7 | +0.28 (padj=0.0034) | ns | ns | ns | ns | +0.29 (padj=0.021) | +0.37 (padj=0.0032) | ns | ns |
| **Homologous recombination** | *rad50* | FBgn0034728 | Not hypoxia-related (discussed in text) | KEGG dme03440 | 31.2 | +0.27 (padj=0.00019) | ns | ns | +0.25 (padj=0.01) | ns | ns | +0.28 (padj=0.0053) | ns | ns |

| | | | | | | | | | | | | | | |
|---|---|---|---|---|---|---|---|---|---|---|---|---|---|---|
| **Homologous recombination** | *okra (Rad54)* | FBgn0002989 | Not hypoxia-related (discussed in text) | KEGG dme03440 | 13 | +0.23 (padj=0.025) | ns | ns | ns | ns | ns | ns | ns | ns |
| **Homologous recombination** | *mus101* | FBgn0002878 | Not hypoxia-related (discussed in text) | KEGG dme03440 | 19.9 | +0.21 (padj=0.042) | ns | ns | ns | ns | ns | ns | ns | ns |
| **Homologous recombination** | *mms4* | FBgn0033549 | Not hypoxia-related (discussed in text) | KEGG dme03440 | 18 | +0.29 (padj=0.048) | ns | ns | ns | ns | ns | ns | ns | ns |
| **Homologous recombination** | *Pol32* | FBgn0027559 | Not hypoxia-related (discussed in text) | KEGG dme03440 | 36.2 | +0.32 (padj=0.0036) | ns | ns | ns | ns | ns | +0.43 (padj=0.00062) | ns | ns |
| **Homologous recombination** | *RpA-70* | FBgn0010173 | Not hypoxia-related (discussed in text) | KEGG dme03440 | 136.1 | +0.21 (padj=0.031) | ns | ns | ns | ns | ns | +0.27 (padj=0.0066) | ns | ns |

**Table S10.** KEGG pathways by treatment comparison and sampling day. KEGG pathways significantly enriched (Benjamini–Hochberg adjusted P < 0.05; clusterProfiler, performed by Novogene) among genes with higher or lower expression in the first-named treatment, pooled across days ("All DEGs") and for each sampling day.

| Comparison | Day | Direction | KEGG_ID | Pathway | Genes | GeneRatio | padj |
|---|---|---|---|---|---|---|---|
| 21 kPa vs Control | Day 2 | Higher | dme04512 | ECM-receptor interaction | 8 | 8/299 | 7.08E-05 |
| 21 kPa vs Control | Day 2 | Higher | dme02010 | ABC transporters | 13 | 13/299 | 0.00101 |
| 21 kPa vs Control | Day 2 | Higher | dme00670 | One carbon pool by folate | 9 | 9/299 | 0.00884 |
| 21 kPa vs Control | Day 2 | Higher | dme03008 | Ribosome biogenesis in eukaryotes | 16 | 16/299 | 0.00921 |
| 21 kPa vs Control | Day 2 | Lower | dme03010 | Ribosome | 51 | 51/322 | 6.07E-17 |
| 21 kPa vs Control | Day 2 | Lower | dme00190 | Oxidative phosphorylation | 37 | 37/322 | 1.89E-08 |
| 21 kPa vs Control | Day 2 | Lower | dme03050 | Proteasome | 18 | 18/322 | 1.21E-06 |
| 21 kPa vs Control | Day 2 | Lower | dme03040 | Spliceosome | 25 | 25/322 | 0.00486 |
| 8 kPa vs 21 kPa | Day 2 | Higher | dme03082 | ATP-dependent chromatin remodeling | 33 | 33/436 | 2.35E-10 |
| 8 kPa vs 21 kPa | Day 2 | Higher | dme03030 | DNA replication | 22 | 22/436 | 2.35E-10 |
| 8 kPa vs 21 kPa | Day 2 | Higher | dme03430 | Mismatch repair | 13 | 13/436 | 1.63E-06 |
| 8 kPa vs 21 kPa | Day 2 | Higher | dme03420 | Nucleotide excision repair | 20 | 20/436 | 2.65E-05 |
| 8 kPa vs 21 kPa | Day 2 | Higher | dme03440 | Homologous recombination | 12 | 12/436 | 0.000335 |
| 8 kPa vs 21 kPa | Day 2 | Higher | dme03040 | Spliceosome | 33 | 33/436 | 0.000335 |
| 8 kPa vs 21 kPa | Day 2 | Higher | dme03022 | Basal transcription factors | 15 | 15/436 | 0.0015 |
| 8 kPa vs 21 kPa | Day 2 | Higher | dme04120 | Ubiquitin mediated proteolysis | 25 | 25/436 | 0.00365 |
| 8 kPa vs 21 kPa | Day 2 | Higher | dme03460 | Fanconi anemia pathway | 11 | 11/436 | 0.00426 |
| 8 kPa vs 21 kPa | Day 2 | Higher | dme04620 | Toll-like receptor signaling pathway | 13 | 13/436 | 0.00633 |
| 8 kPa vs 21 kPa | Day 2 | Higher | dme03410 | Base excision repair | 11 | 11/436 | 0.00998 |
| 8 kPa vs 21 kPa | Day 2 | Higher | dme04068 | FoxO signaling pathway | 17 | 17/436 | 0.0105 |
| 8 kPa vs 21 kPa | Day 2 | Higher | dme04310 | Wnt signaling pathway | 22 | 22/436 | 0.0105 |
| 8 kPa vs 21 kPa | Day 2 | Higher | dme03013 | Nucleocytoplasmic transport | 21 | 21/436 | 0.0411 |
| 8 kPa vs 21 kPa | Day 2 | Lower | dme01200 | Carbon metabolism | 49 | 49/523 | 5.37E-12 |

| | | | | | | | |
|---|---|---|---|---|---|---|---|
| 8 kPa vs 21 kPa | Day 2 | Lower | dme01210 | 2-Oxocarboxylic acid metabolism | 23 | 23/523 | 1.48E-10 |
| 8 kPa vs 21 kPa | Day 2 | Lower | dme00020 | Citrate cycle (TCA cycle) | 23 | 23/523 | 7.30E-10 |
| 8 kPa vs 21 kPa | Day 2 | Lower | dme04820 | Cytoskeleton in muscle cells | 29 | 29/523 | 1.34E-09 |
| 8 kPa vs 21 kPa | Day 2 | Lower | dme00670 | One carbon pool by folate | 18 | 18/523 | 1.52E-08 |
| 8 kPa vs 21 kPa | Day 2 | Lower | dme04512 | ECM-receptor interaction | 11 | 11/523 | 3.33E-08 |
| 8 kPa vs 21 kPa | Day 2 | Lower | dme01230 | Biosynthesis of amino acids | 29 | 29/523 | 1.77E-07 |
| 8 kPa vs 21 kPa | Day 2 | Lower | dme00260 | Glycine, serine and threonine metabolism | 16 | 16/523 | 1.19E-06 |
| 8 kPa vs 21 kPa | Day 2 | Lower | dme00630 | Glyoxylate and dicarboxylate metabolism | 16 | 16/523 | 9.77E-06 |
| 8 kPa vs 21 kPa | Day 2 | Lower | dme00785 | Lipoic acid metabolism | 11 | 11/523 | 9.66E-05 |
| 8 kPa vs 21 kPa | Day 2 | Lower | dme00270 | Cysteine and methionine metabolism | 17 | 17/523 | 0.000116 |
| 8 kPa vs 21 kPa | Day 2 | Lower | dme00010 | Glycolysis / Gluconeogenesis | 20 | 20/523 | 0.000185 |
| 8 kPa vs 21 kPa | Day 2 | Lower | dme00250 | Alanine, aspartate and glutamate metabolism | 15 | 15/523 | 0.000364 |
| 8 kPa vs 21 kPa | Day 2 | Lower | dme04142 | Lysosome | 35 | 35/523 | 0.00047 |
| 8 kPa vs 21 kPa | Day 2 | Lower | dme00030 | Pentose phosphate pathway | 11 | 11/523 | 0.000548 |
| 8 kPa vs 21 kPa | Day 2 | Lower | dme02010 | ABC transporters | 16 | 16/523 | 0.00082 |
| 8 kPa vs 21 kPa | Day 2 | Lower | dme04981 | Folate transport and metabolism | 11 | 11/523 | 0.000869 |
| 8 kPa vs 21 kPa | Day 2 | Lower | dme00640 | Propanoate metabolism | 13 | 13/523 | 0.000977 |
| 8 kPa vs 21 kPa | Day 2 | Lower | dme00220 | Arginine biosynthesis | 10 | 10/523 | 0.00152 |
| 8 kPa vs 21 kPa | Day 2 | Lower | dme00280 | Valine, leucine and isoleucine degradation | 14 | 14/523 | 0.0016 |
| 8 kPa vs 21 kPa | Day 2 | Lower | dme00620 | Pyruvate metabolism | 16 | 16/523 | 0.0018 |
| 8 kPa vs 21 kPa | Day 2 | Lower | dme00380 | Tryptophan metabolism | 10 | 10/523 | 0.00367 |
| 8 kPa vs 21 kPa | Day 2 | Lower | dme01212 | Fatty acid metabolism | 17 | 17/523 | 0.0088 |
| 8 kPa vs 21 kPa | Day 2 | Lower | dme00310 | Lysine degradation | 13 | 13/523 | 0.00968 |
| 8 kPa vs 21 kPa | Day 2 | Lower | dme04145 | Phagosome | 21 | 21/523 | 0.0133 |
| 8 kPa vs 21 kPa | Day 2 | Lower | dme00230 | Purine metabolism | 24 | 24/523 | 0.0133 |
| 8 kPa vs 21 kPa | Day 2 | Lower | dme00410 | beta-Alanine metabolism | 10 | 10/523 | 0.021 |
| 8 kPa vs 21 kPa | Day 2 | Lower | dme00040 | Pentose and glucuronate interconversions | 15 | 15/523 | 0.0325 |

| | | | | | | | |
|---|---|---|---|---|---|---|---|
| 8 kPa vs 21 kPa | Day 2 | Lower | dme00 511 | Other glycan degradation | 8 | 8/523 | 0.0353 |
| 8 kPa vs 21 kPa | Day 2 | Lower | dme00 061 | Fatty acid biosynthesis | 6 | 6/523 | 0.0402 |
| 8 kPa vs 21 kPa | Day 2 | Lower | dme01 240 | Biosynthesis of cofactors | 33 | 33/523 | 0.0411 |
| 8 kPa vs Control | Day 2 | Higher | dme03 030 | DNA replication | 21 | 21/197 | 4.03E-16 |
| 8 kPa vs Control | Day 2 | Higher | dme03 410 | Base excision repair | 11 | 11/197 | 2.57E-05 |
| 8 kPa vs Control | Day 2 | Higher | dme03 420 | Nucleotide excision repair | 13 | 13/197 | 6.45E-05 |
| 8 kPa vs Control | Day 2 | Higher | dme03 440 | Homologous recombination | 9 | 9/197 | 0.0001 11 |
| 8 kPa vs Control | Day 2 | Higher | dme03 430 | Mismatch repair | 8 | 8/197 | 0.0001 11 |
| 8 kPa vs Control | Day 2 | Higher | dme03 082 | ATP-dependent chromatin remodeling | 14 | 14/197 | 0.0006 44 |
| 8 kPa vs Control | Day 2 | Higher | dme03 460 | Fanconi anemia pathway | 8 | 8/197 | 0.0015 3 |
| 8 kPa vs Control | Day 2 | Higher | dme03 013 | Nucleocytoplasmic transport | 14 | 14/197 | 0.0080 1 |
| 8 kPa vs Control | Day 2 | Higher | dme03 083 | Polycomb repressive complex | 9 | 9/197 | 0.0093 8 |
| 8 kPa vs Control | Day 2 | Higher | dme04 350 | TGF-beta signaling pathway | 9 | 9/197 | 0.01 |
| 8 kPa vs Control | Day 2 | Lower | dme03 010 | Ribosome | 61 | 61/301 | 1.09E-27 |
| 8 kPa vs Control | Day 2 | Lower | dme00 190 | Oxidative phosphorylation | 49 | 49/301 | 1.30E-18 |
| 8 kPa vs Control | Day 2 | Lower | dme00 020 | Citrate cycle (TCA cycle) | 11 | 11/301 | 0.0040 1 |
| 8 kPa vs Control | Day 2 | Lower | dme00 620 | Pyruvate metabolism | 12 | 12/301 | 0.0060 1 |
| 8 kPa vs Control | Day 2 | Lower | dme01 210 | 2-Oxocarboxylic acid metabolism | 10 | 10/301 | 0.0067 3 |
| 8 kPa vs Control | Day 2 | Lower | dme00 010 | Glycolysis / Gluconeogenesis | 12 | 12/301 | 0.0236 |
| 21 kPa vs Control | Day 6 | Higher | dme03 030 | DNA replication | 20 | 20/149 | 4.66E-17 |
| 21 kPa vs Control | Day 6 | Higher | dme03 430 | Mismatch repair | 6 | 6/149 | 0.0071 9 |
| 21 kPa vs Control | Day 6 | Higher | dme03 410 | Base excision repair | 7 | 7/149 | 0.0108 |
| 21 kPa vs Control | Day 6 | Higher | dme03 420 | Nucleotide excision repair | 8 | 8/149 | 0.0335 |
| 21 kPa vs Control | Day 6 | Lower | dme03 010 | Ribosome | 67 | 67/256 | 1.05E-38 |
| 21 kPa vs Control | Day 6 | Lower | dme04 140 | Autophagy - animal | 20 | 20/256 | 0.0206 |
| 8 kPa vs 21 kPa | Day 6 | Higher | dme03 030 | DNA replication | 9 | 9/189 | 0.0106 |
| 8 kPa vs 21 kPa | Day 6 | Lower | dme03 010 | Ribosome | 68 | 68/237 | 1.98E-42 |

| | | | | | | | |
|---|---|---|---|---|---|---|---|
| 8 kPa vs 21 kPa | Day 6 | Lower | dme00190 | Oxidative phosphorylation | 40 | 40/237 | 1.92E-15 |
| 8 kPa vs Control | Day 6 | Higher | dme03030 | DNA replication | 28 | 28/431 | 1.16E-17 |
| 8 kPa vs Control | Day 6 | Higher | dme03410 | Base excision repair | 18 | 18/431 | 1.16E-07 |
| 8 kPa vs Control | Day 6 | Higher | dme03430 | Mismatch repair | 14 | 14/431 | 1.16E-07 |
| 8 kPa vs Control | Day 6 | Higher | dme03420 | Nucleotide excision repair | 21 | 21/431 | 5.70E-06 |
| 8 kPa vs Control | Day 6 | Higher | dme03440 | Homologous recombination | 12 | 12/431 | 0.000392 |
| 8 kPa vs Control | Day 6 | Higher | dme03460 | Fanconi anemia pathway | 12 | 12/431 | 0.00143 |
| 8 kPa vs Control | Day 6 | Higher | dme03083 | Polycomb repressive complex | 15 | 15/431 | 0.0058 |
| 8 kPa vs Control | Day 6 | Higher | dme03082 | ATP-dependent chromatin remodeling | 20 | 20/431 | 0.00661 |
| 8 kPa vs Control | Day 6 | Higher | dme03013 | Nucleocytoplasmic transport | 23 | 23/431 | 0.0136 |
| 8 kPa vs Control | Day 6 | Lower | dme03010 | Ribosome | 89 | 89/484 | 1.02E-41 |
| 8 kPa vs Control | Day 6 | Lower | dme00190 | Oxidative phosphorylation | 52 | 52/484 | 7.64E-12 |
| 8 kPa vs Control | Day 6 | Lower | dme00260 | Glycine, serine and threonine metabolism | 11 | 11/484 | 0.0286 |
| 21 kPa vs Control | Pooled | All DEGs | dme03010 | Ribosome | 65 | 65/462 | 6.81E-20 |
| 21 kPa vs Control | Pooled | All DEGs | dme00190 | Oxidative phosphorylation | 44 | 44/462 | 1.13E-07 |
| 8 kPa vs Control | Pooled | All DEGs | dme03010 | Ribosome | 97 | 97/935 | 1.69E-24 |
| 8 kPa vs Control | Pooled | All DEGs | dme03030 | DNA replication | 30 | 30/935 | 2.44E-11 |
| 8 kPa vs Control | Pooled | All DEGs | dme00190 | Oxidative phosphorylation | 69 | 69/935 | 9.59E-09 |
| 8 kPa vs Control | Pooled | All DEGs | dme03430 | Mismatch repair | 16 | 16/935 | 2.02E-05 |
| 8 kPa vs Control | Pooled | All DEGs | dme03440 | Homologous recombination | 16 | 16/935 | 0.00267 |
| 8 kPa vs Control | Pooled | All DEGs | dme03410 | Base excision repair | 18 | 18/935 | 0.00596 |

**Table S11.** Median and interquartile range of DESeq2 log2 fold changes across all genes in four KEGG pathways (ribosome, dme03010; oxidative phosphorylation, dme00190; glycolysis/gluconeogenesis, dme00010; citrate cycle, dme00020), with the number of pathway genes significantly higher or lower (adjusted P < 0.05). Fold changes are for the first-named treatment; no additional tests were performed. 8 kPa = Chamber PO2 ≈ 8 kPa; 21 kPa = Chamber PO2 ≈ 21 kPa; Control = outside-chamber PO2 ≈ 21 kPa.

| Comparison | Day | Pathway | KEGG_ID | Genes_tested | Median_log2FC | IQR | Higher_padj05 | Lower_padj05 |
|---|---|---|---|---|---|---|---|---|
| 21 kPa vs Control | Pooled | Citrate cycle (TCA cycle) | dme00020 | 31 | 0.034 | -0.04 to 0.07 | 0 | 0 |
| 8 kPa vs 21 kPa | Pooled | Citrate cycle (TCA cycle) | dme00020 | 33 | -0.094 | -0.16 to -0.06 | 0 | 0 |
| 8 kPa vs Control | Pooled | Citrate cycle (TCA cycle) | dme00020 | 32 | -0.062 | -0.12 to 0.01 | 0 | 3 |
| 21 kPa vs Control | Day 2 | Citrate cycle (TCA cycle) | dme00020 | 30 | 0.095 | 0.03 to 0.17 | 3 | 0 |
| 8 kPa vs 21 kPa | Day 2 | Citrate cycle (TCA cycle) | dme00020 | 31 | -0.204 | -0.3 to -0.14 | 0 | 17 |
| 8 kPa vs Control | Day 2 | Citrate cycle (TCA cycle) | dme00020 | 30 | -0.11 | -0.16 to -0.05 | 0 | 2 |
| 21 kPa vs Control | Day 6 | Citrate cycle (TCA cycle) | dme00020 | 29 | -0.034 | -0.12 to 0 | 0 | 1 |
| 8 kPa vs 21 kPa | Day 6 | Citrate cycle (TCA cycle) | dme00020 | 30 | 0.033 | -0.03 to 0.1 | 0 | 0 |
| 8 kPa vs Control | Day 6 | Citrate cycle (TCA cycle) | dme00020 | 30 | -0.029 | -0.1 to 0.05 | 1 | 4 |
| 21 kPa vs Control | Pooled | Glycolysis / Gluconeogenesis | dme00010 | 49 | 0.022 | -0.09 to 0.24 | 0 | 0 |
| 8 kPa vs 21 kPa | Pooled | Glycolysis / Gluconeogenesis | dme00010 | 48 | -0.142 | -0.36 to 0.04 | 0 | 0 |
| 8 kPa vs Control | Pooled | Glycolysis / Gluconeogenesis | dme00010 | 48 | -0.077 | -0.24 to 0.04 | 1 | 4 |
| 21 kPa vs Control | Day 2 | Glycolysis / Gluconeogenesis | dme00010 | 46 | 0.038 | -0.1 to 0.34 | 0 | 2 |
| 8 kPa vs 21 kPa | Day 2 | Glycolysis / Gluconeogenesis | dme00010 | 47 | -0.18 | -0.44 to 0.02 | 0 | 13 |
| 8 kPa vs Control | Day 2 | Glycolysis / Gluconeogenesis | dme00010 | 48 | -0.113 | -0.27 to -0.01 | 0 | 4 |
| 21 kPa vs Control | Day 6 | Glycolysis / Gluconeogenesis | dme00010 | 47 | -0.046 | -0.18 to 0.06 | 0 | 0 |
| 8 kPa vs 21 kPa | Day 6 | Glycolysis / Gluconeogenesis | dme00010 | 47 | -0.002 | -0.11 to 0.06 | 0 | 0 |
| 8 kPa vs Control | Day 6 | Glycolysis / Gluconeogenesis | dme00010 | 45 | -0.065 | -0.35 to 0.08 | 2 | 3 |
| 21 kPa vs Control | Pooled | Oxidative phosphorylation | dme00190 | 121 | -0.053 | -0.18 to 0.07 | 9 | 9 |

| | | | | | | | | |
|---|---|---|---|---|---|---|---|---|
| 8 kPa vs 21 kPa | Pooled | Oxidative phosphorylation | dme00 190 | 122 | -0.101 | -0.15 to - 0.05 | 0 | 0 |
| 8 kPa vs Control | Pooled | Oxidative phosphorylation | dme00 190 | 120 | -0.146 | -0.32 to - 0.02 | 8 | 41 |
| 21 kPa vs Control | Day 2 | Oxidative phosphorylation | dme00 190 | 120 | -0.04 | -0.26 to 0.14 | 12 | 19 |
| 8 kPa vs 21 kPa | Day 2 | Oxidative phosphorylation | dme00 190 | 121 | -0.076 | -0.2 to 0.07 | 6 | 20 |
| 8 kPa vs Control | Day 2 | Oxidative phosphorylation | dme00 190 | 119 | -0.151 | -0.25 to 0 | 7 | 23 |
| 21 kPa vs Control | Day 6 | Oxidative phosphorylation | dme00 190 | 117 | -0.035 | -0.13 to 0.04 | 4 | 0 |
| 8 kPa vs 21 kPa | Day 6 | Oxidative phosphorylation | dme00 190 | 117 | -0.132 | -0.28 to - 0.01 | 0 | 6 |
| 8 kPa vs Control | Day 6 | Oxidative phosphorylation | dme00 190 | 118 | -0.145 | -0.38 to 0 | 6 | 43 |
| 21 kPa vs Control | Pooled | Ribosome | dme03 010 | 169 | -0.108 | -0.22 to - 0.03 | 1 | 36 |
| 8 kPa vs 21 kPa | Pooled | Ribosome | dme03 010 | 169 | -0.095 | -0.13 to - 0.02 | 0 | 0 |
| 8 kPa vs Control | Pooled | Ribosome | dme03 010 | 167 | -0.192 | -0.32 to - 0.08 | 2 | 81 |
| 21 kPa vs Control | Day 2 | Ribosome | dme03 010 | 167 | -0.123 | -0.25 to 0 | 1 | 32 |
| 8 kPa vs 21 kPa | Day 2 | Ribosome | dme03 010 | 168 | 0.009 | -0.07 to 0.1 | 9 | 8 |
| 8 kPa vs Control | Day 2 | Ribosome | dme03 010 | 166 | -0.122 | -0.19 to - 0.04 | 1 | 35 |
| 21 kPa vs Control | Day 6 | Ribosome | dme03 010 | 166 | -0.125 | -0.22 to - 0.02 | 0 | 13 |
| 8 kPa vs 21 kPa | Day 6 | Ribosome | dme03 010 | 166 | -0.175 | -0.26 to - 0.06 | 0 | 16 |
| 8 kPa vs Control | Day 6 | Ribosome | dme03 010 | 166 | -0.279 | -0.46 to -0.1 | 0 | 90 |

### Supplementary Figures

**Figure S1 Modified recombination testcross.** Crossing schemes for a) the marker-free control crosse and b) the multiply-marked crosses on identical genetic backgrounds.

a

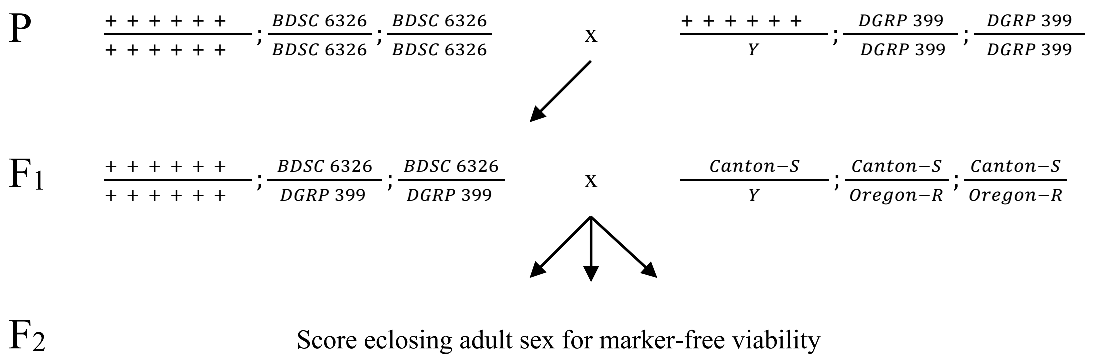


b

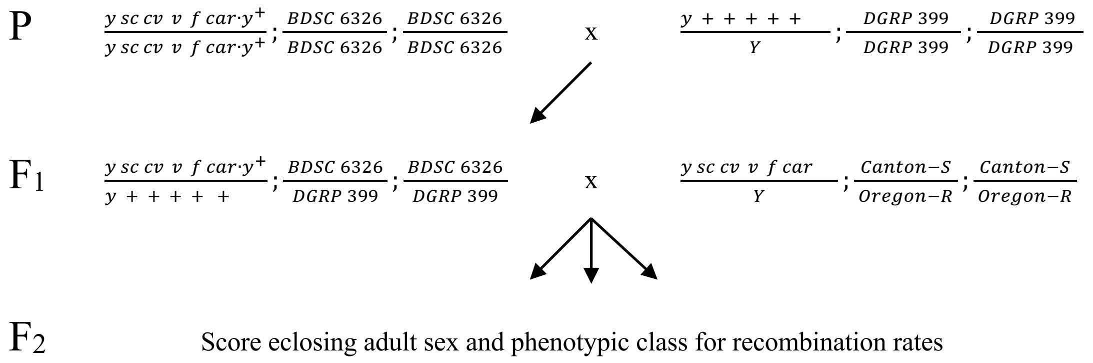

**Figure S2 Digital egg counts and egg density.** (a) Digital egg-count standard curves: digital egg counts from two observers (CH, EH) plotted against microscope counts of the same 30 vials (calibration dataset of Schneider et al. 2025). Solid lines are Type II (major axis) regressions; the dashed line is 1:1. Digital counts were corrected as (digital count − intercept)/slope. (b) Angular-transformed $F_2$ egg-to-adult viability against eggs per vial for marker-free and multiply-marked crosses. Points are vials by $F_2$ sex; lines are least-squares fits with 95% confidence bands.

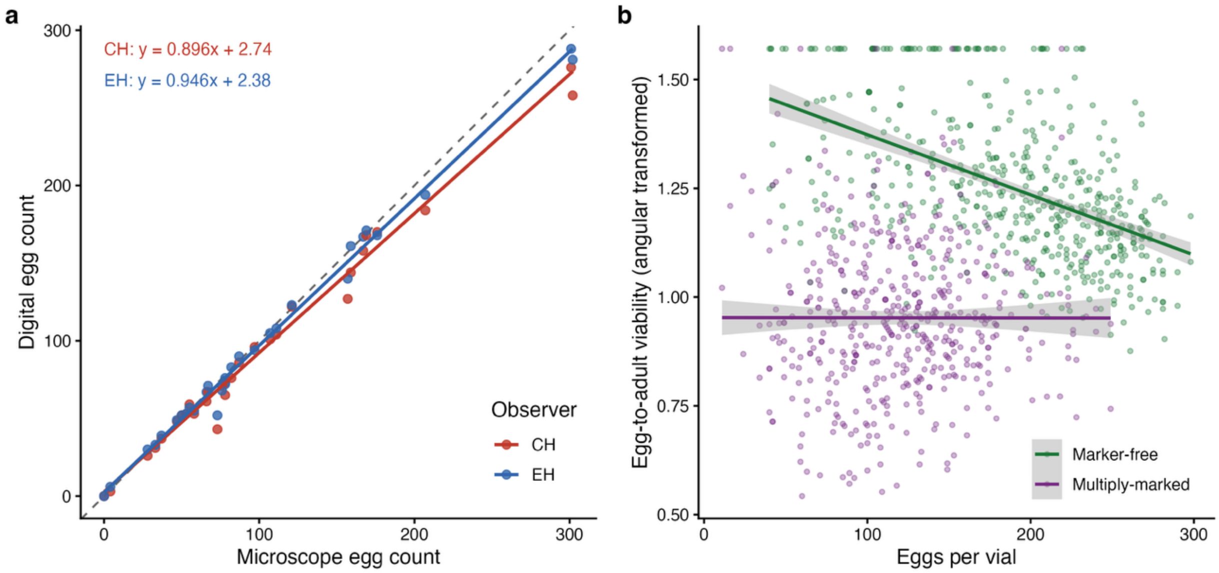

**Figure S3 Differential gene expression between treatments across two time points.** This figure complements Figure 2 in the main text by showing the same treatment comparisons separately for each sampling day, using four independent biological replicates of ovaries from marker-free females. (a–c) Day 2 and (d–f) Day 6: (a, d) Chamber $PO_2$ 8 kPa vs. Control; (b, e) Chamber $PO_2$ 21 kPa vs. Control; (c, f) Chamber $PO_2$ 21 kPa vs. Chamber $PO_2$ 8 kPa. Each point is a gene, plotted by $\log_2$ fold change and adjusted P-value; colors indicate the treatment with higher expression (adjusted $P < 0.05$), and the five most significant genes on each side are labeled. (g) Overlap of DEGs among the three comparisons, computed separately for Day 2 (light blue) and Day 6 (dark blue). Connected dots indicate genes differentially expressed in that combination of comparisons; vertical bars give the number of genes in each intersection on each day, and horizontal bars give the total number of DEGs per comparison (table S8).

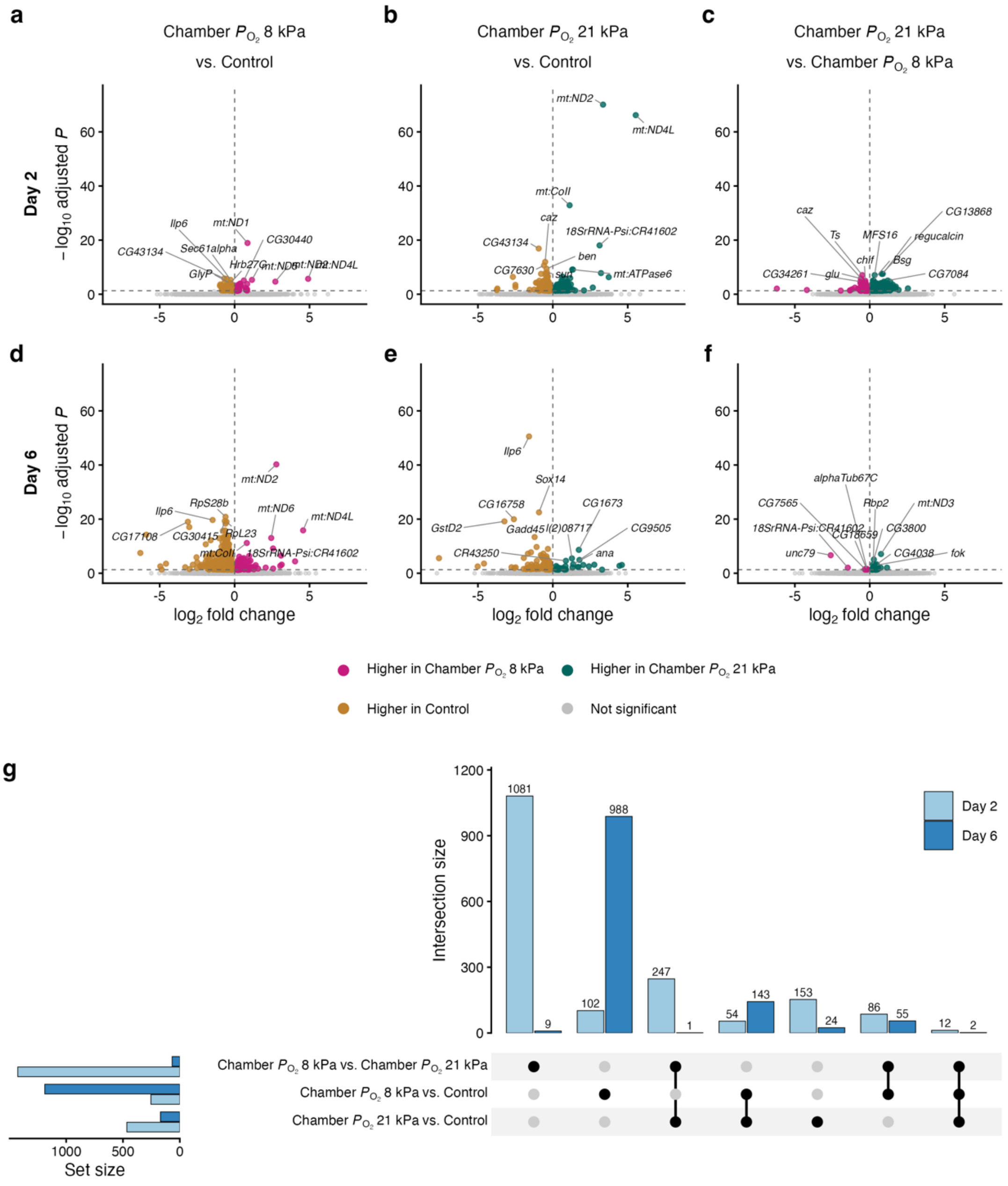

**Figure S4 Gene Ontology enrichment among upregulated genes across treatments.** Dot plots show GO terms significantly enriched (adjusted $p < 0.05$) among genes upregulated in Chamber $P_{O2}$ 21 kPa vs. Control (left) and Chamber $P_{O2}$ 8 kPa vs. Control (right), across both time points. Upregulated genes are those with higher expression in the chamber treatment than in the outside-chamber control. The x-axis shows the gene ratio, dot size indicates the number of genes associated with each term, and dot color indicates enrichment significance as $-\log_{10}$(adjusted p-value). Terms are ordered by their maximum enrichment significance across comparisons.

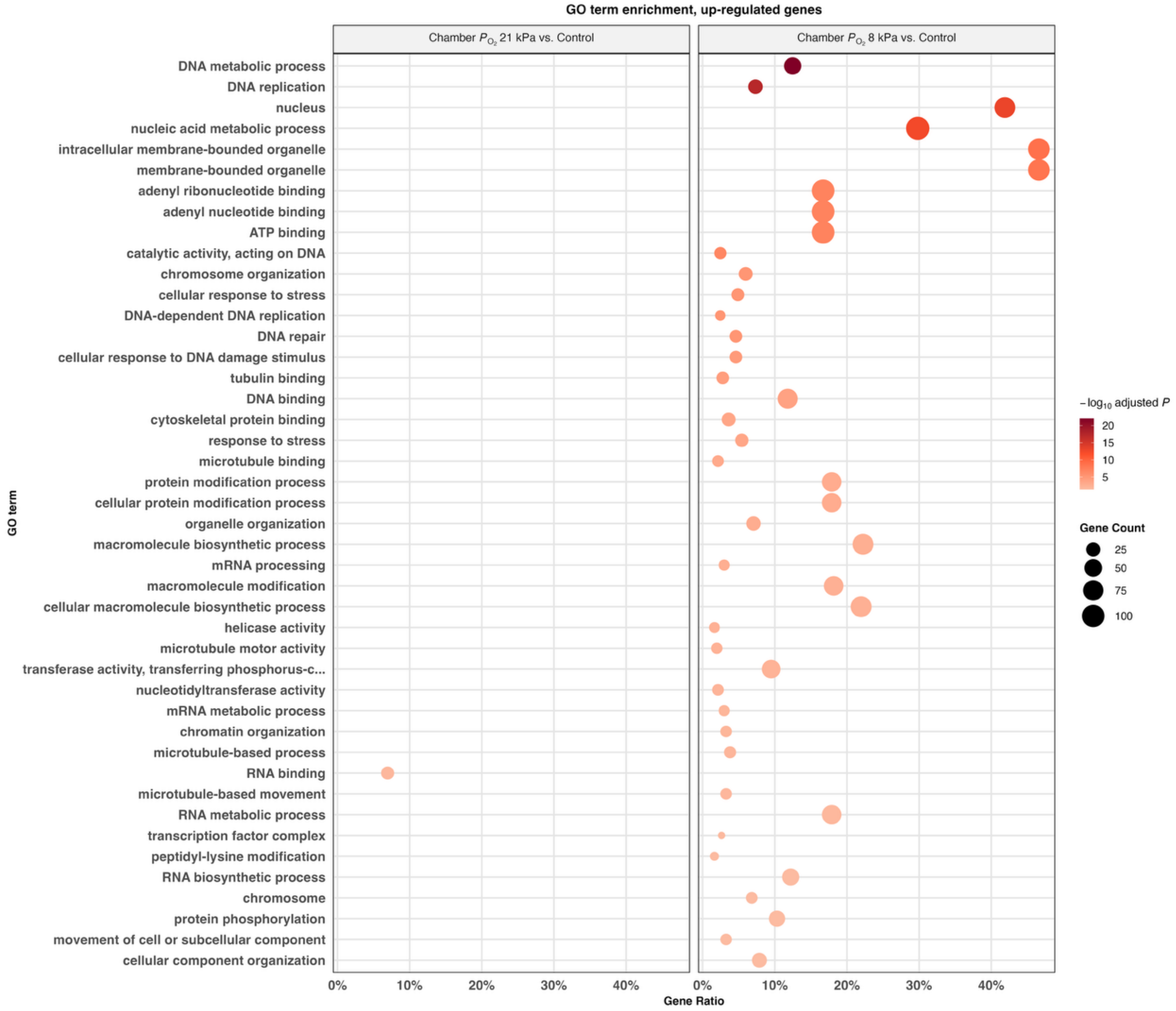

**Figure S5. Gene Ontology enrichment among downregulated genes across treatments.** Dot plots show GO terms significantly enriched (adjusted $p < 0.05$) among genes downregulated in Chamber $P_{O2}$ 21 kPa vs. Control (left) and Chamber $P_{O2}$ 8 kPa vs. Control (right), across both time points. Downregulated genes are those with lower expression in the chamber treatment than in the outside-chamber control. The x-axis shows the gene ratio, dot size indicates the number of genes associated with each term, and dot color indicates enrichment significance as $-\log_{10}$(adjusted p-value). Terms are ordered by their maximum enrichment significance across comparisons.

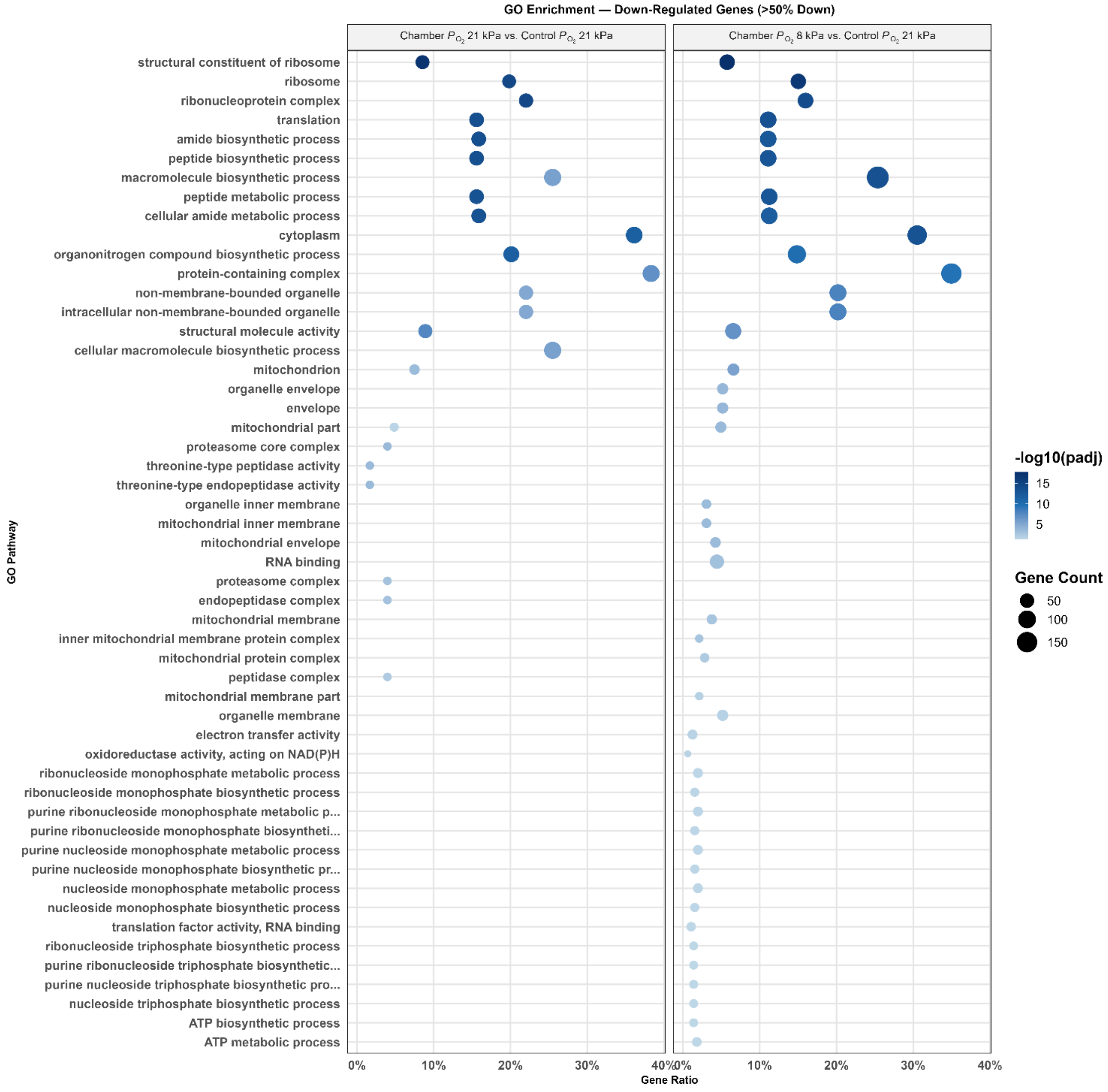

**Figure S6. Differential gene expression of core HIF alpha pathway genes.** Ovarian expression of core HIF pathway genes. Expression (FPKM) of (a) *sima* (HIF-α), (b) *tango* (tgo; HIF-β/ARNT), (c) *fatiga* (fga; HIF prolyl hydroxylase), and (d) *Vhl* (von Hippel-Lindau) in $F_1$ ovaries 2 and 6 days after eclosion, under Control PO2 ≈ 21 kPa (gold), Chamber PO2 ≈ 21 kPa (teal), and Chamber PO2 ≈ 8 kPa (pink). Large points and bars show the mean ± SD of four biological replicates; small points are individual replicates. The text above each panel gives the log2 fold change and Benjamini–Hochberg adjusted P for the pooled Chamber PO2 ≈ 8 kPa vs Control comparison (Chamber PO2 ≈ 8 kPa vs Chamber PO2 ≈ 21 kPa: *fatiga* log2 fold change 0.13, not significant; Table S9) from the genome-wide DESeq2 analysis (* adjusted $P < 0.05$); no additional tests were performed. Results for all pairwise comparisons, pooled and by day, are given in Table S9.

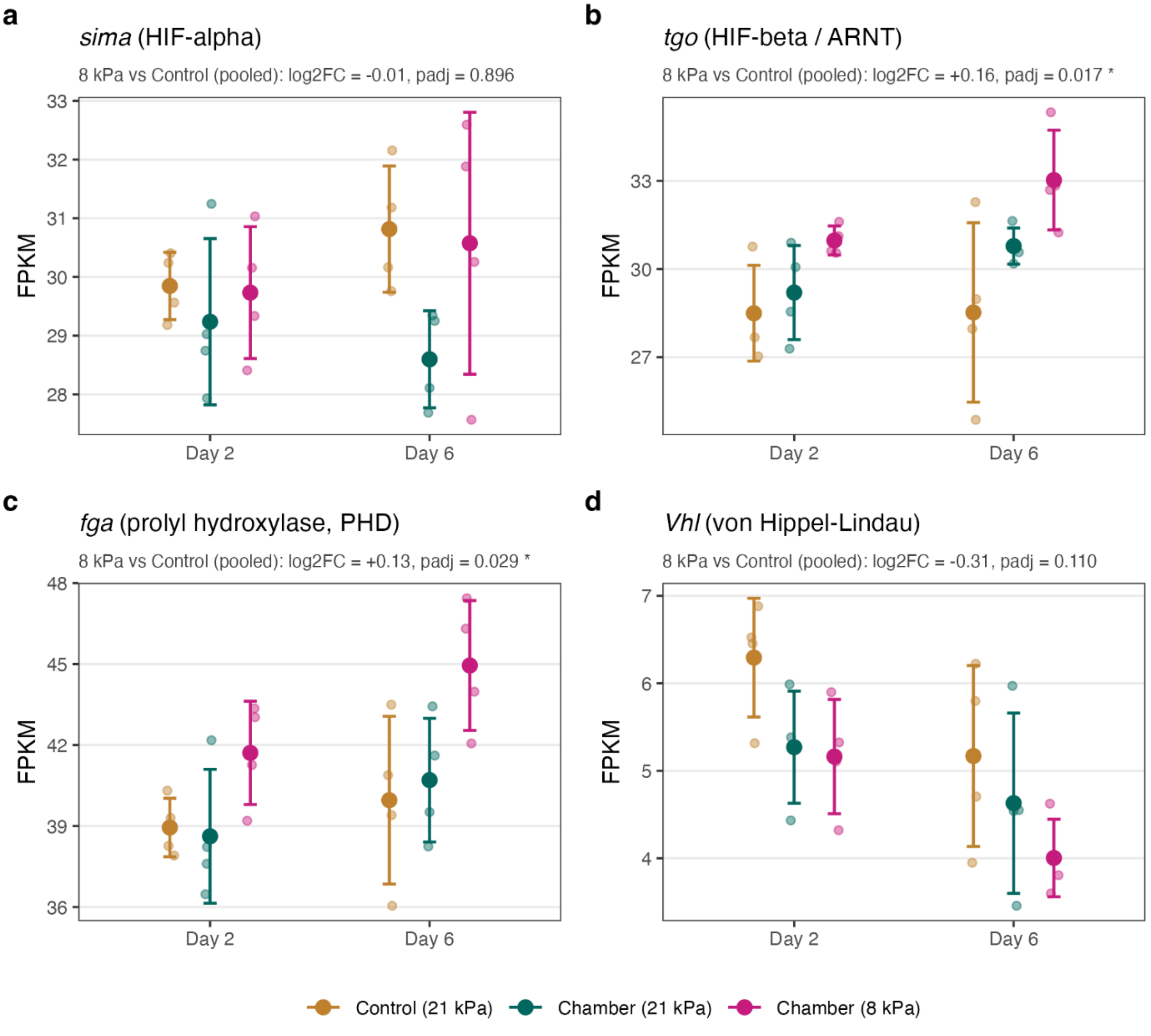
